# Motor Protein MYO1C is Critical for Photoreceptor Opsin Trafficking and Visual Function

**DOI:** 10.1101/2020.06.02.129890

**Authors:** Ashish K. Solanki, Stephen Walterhouse, René Martin, Elisabeth Obert, Ehtesham Arif, Bushra Rahman, Barbel Rohrer, Joshua Lipschutz, Rupak D. Mukherjee, Russell A. Norris, Jeffery Sundstrom, Hans-Joachim Knölker, Shahid Husain, Manas R. Biswal, Deepak Nihalani, Glenn P. Lobo

## Abstract

Unconventional myosins linked to deafness are also proposed to play a role in retinal cell physiology. However, their direct role in photoreceptor function remains unclear. We demonstrate that systemic loss of the unconventional myosin MYO1C in mice specifically affected opsin trafficking, leading to loss of visual function. Electroretinogram analysis of *Myo1c* knockout (*Myo1c*-KO) mice showed a progressive loss of photoreceptor function. Immunohistochemistry and binding assays demonstrated MYO1C localization to photoreceptor inner and outer segments (OS) and identified a direct interaction of rhodopsin with the MYO1C cargo domain. In *Myo1c*-KO retinas, rhodopsin mislocalized to rod inner segments (IS) and cell bodies, while cone opsins in OS showed punctate staining. In aged mice, the histological and ultrastructural examination of the phenotype of *Myo1c*-KO retinas showed progressively shorter photoreceptor OS. These results demonstrate that MYO1C is critical for opsin trafficking to the photoreceptor OS and for normal visual function.

## Introduction

Protein trafficking within the photoreceptors must occur efficiently and at high fidelity for photoreception, photoreceptor structural maintenance, and overall retinal cell homeostasis. Additionally, it is well-known that proper opsin trafficking is tightly coupled to photoreceptor cell survival and function [1-9]. However, the cellular events that participate in retinal injuries due to improper signaling and protein trafficking to the photoreceptor outer segments (OS), are not yet fully understood. While many proteins are known to play an essential role in retinal cell development and function, the involvement of motor proteins in eye biology is less understood. Identification of genetic mutations in the *Myo7a* gene associated with retinal degeneration in Usher syndrome suggests that unconventional myosins play a critical role in retinal pigmented epithelium (RPE) and photoreceptor cell function [10, 11]. Unconventional myosins are motor proteins that are proposed to transport membranous organelles along the actin filaments in an adenosine triphosphate (ATP)-dependent manner, and additional roles are currently being discovered [12-14]. The loss of *Myo7a* primarily affects RPE and OS phagocytosis leading to retinal cell degeneration. However, it is believed that other yet unidentified class I myosins may participate more directly in photoreceptor cell function. Here we present compelling evidence for another unconventional actin-binding motor protein, MYO1C, with its primary localization to photoreceptors that plays an important role in retinal cell structure and function via opsin trafficking to the photoreceptor OS.

Rhodopsin and cone pigments in photoreceptor OS mediate scotopic and photopic vision, respectively. The visual pigment rhodopsin is a prototypical G-protein coupled receptor (GPCR) expressed by retinal rods for photon absorption. Light sensitivity is conferred by 11-*cis* retinaldehyde, a chromophore that is covalently linked to the K296 residue of the opsin protein [15-19] Photon absorption causes a *cis* to *trans* conformational shift in the retinaldehyde, leading to structural changes in the opsin protein moiety [6]. This initiates a GPCR signaling pathway/phototransduction cascade, signaling the presence of light. Each photoreceptor cell contains an OS housing the phototransduction machinery, an inner segment (IS) where proteins are biosynthesized, and a synaptic terminal for signal transmission. One of the fundamental steps in vision is the proper assembly of signal-transducing membranes, including the transport and sorting of protein components. A major cause of neurodegenerative and other inherited retinal disorders is the improper localization of proteins. Mislocalization of the dim-light photoreceptor protein rhodopsin is a phenotype observed in many forms of blinding diseases, including retinitis pigmentosa (RP). The proteins that participate in phototransduction (including rhodopsin, transducin, phosphodiesterase [PDE6], or the cyclic nucleotide-gated channels [CNG]) are synthesized in the IS and must be transported through the connecting cilium to the OS. These proteins are either transmembrane or peripherally associated membrane, which are attached to the membrane surface [1-9]. How the transmembrane proteins (e.g., rhodopsin and CNG) and peripherally associated proteins (e.g., transducin and PDE6) traffic through the IS to incorporate eventually in the nascent disc membrane or the photoreceptor outer membrane is not fully understood and constitutes an area of intense research, as improper trafficking of these protein causes retinal cell degeneration and can lead to blindness [1-9].

Genetic mutations in myosins that lead to hearing loss have also been associated with retinal degeneration. Some of the essential genes involved in either or both of these functions belong to a family of unconventional motor proteins and include MYO3A [20], MYO7A, MYO6, MYO15 [20-22], and MYO5. Recently, it was reported that another unconventional myosin, MYO1C, where mutations affected its nucleotide-binding pocket and calcium binding ability and these were associated with deafness [23]. Importantly, MYO1C was identified in proteomic analysis of the retina and vitreous fluid as part of a protein hub involved in oxidative stress [23-25]. MYO1C is an actin-binding motor protein that is widely expressed in multiple cell types. It participates in a variety of cellular functions, including protein trafficking and translocation [12, 26-28]. As MYO1C has low tissue specificity based on mRNA and protein expression, it remains unclear which cell type is most dependent on MYO1C trafficking function and is affected by the loss of MYO1C.

In this study, we systematically analyzed the function of the unconventional motor protein MYO1C in protein trafficking in photoreceptors. We found that a global genetic deletion of *Myo1c* resulted in a retinal phenotype only, which manifested as a progressive mistrafficking of opsins to the OS. Using retinal lysate from wild-type (WT) mice in co-immunoprecipitation assays, we showed that MYO1C and rhodopsin directly interact, indicating that opsin was a cargo for MYO1C. Loss of MYO1C promoted a progressive shortening of OSs that was concomitant with a reduction in photoreceptor function, suggesting that MYO1C is critical for maintenance of photoreceptor cell structure and for visual function. Our findings have significant clinical implications for degenerative rod and cone diseases, as mutations in MYO1C or its interacting partners are predicted to affect retinal health and visual function by altering opsin trafficking to the photoreceptor OS, a fundamental step for maintaining visual function in humans.

## Results

### Construction and Validation of *Myo1c* Null Mice

We previously generated *Myo1c* floxed mice using the standard knockout strategy [29] (**Fig. S1a**). Systemic deletion of *Myo1c* was achieved by crossing *Myo1c* floxed (*Myo1cfl/fl*) mice with Actin Cre+ (ActCre+ JAX labs) mice to generate *Myo1cfl/fl-ActCre+/-* knockout mice (referred to as *Myo1c*-KO mice in this manuscript). Western blotting of protein lysates from various tissues including kidney, heart, and liver of *Myo1c*-KO mice showed complete loss of MYO1C, thus confirming the systemic deletion of *Myo1c* (**Fig. S1b**). Additionally, immunofluorescence expression analysis of these tissues further confirmed loss of MYO1C protein in *Myo1c*-KO mice (**Figs. S2a-c**).

### Genetic Deletion of *Myo1c* induced Visual Impairment in Mice

Immunofluorescence analysis showed that MYO1C was enriched in the rod photoreceptor outer (OS) and inner segments (IS) (**Fig. 1a**), and also in cone photoreceptor OS of wild-type (WT) mice (**Fig. 1b**), but absent in photoreceptors of *Myo1c*-KO animals (**Figs. 1a** and **1c**). Western blot analysis further confirmed that MYO1C protein was absent in the retinas of *Myo1c*-KO mice (**Fig. 1d**). Since mutations or deletion of the motor protein MYO7A were associated with retinal degeneration in Usher syndrome and its animal model, it prompted us to investigate the effect of *Myo1c* in retinal function. Using electroretinograms (ERGs) [30, 31], we tested photoreceptor cell function of *Myo1c*-KO and WT mice (*n*=8 mice per genotype and age-group; 50:50 ratio of male and female) under dark-adapted scotopic conditions. In contrast to WT animals, we observed reduced ERGs for *Myo1c*-KO mice at different ages. Two month old *Myo1c*-KO mice showed a significant reduction in the ***a***-wave amplitudes, but not in ***b***-wave amplitudes (*p*<0.0068 and *p*<0.098, respectively) (**Figs. 2a** and **2c**). Strikingly, ERG analysis of adult six months old *Myo1c*-KO mice showed severe loss of retinal function, in which a significant reduction in both ***a***- and ***b***-waves was observed (38-45% lower than WT animals (^**^*p*<0.005; **Figs. 2b** and **2d)**.

**Fig. 1:**
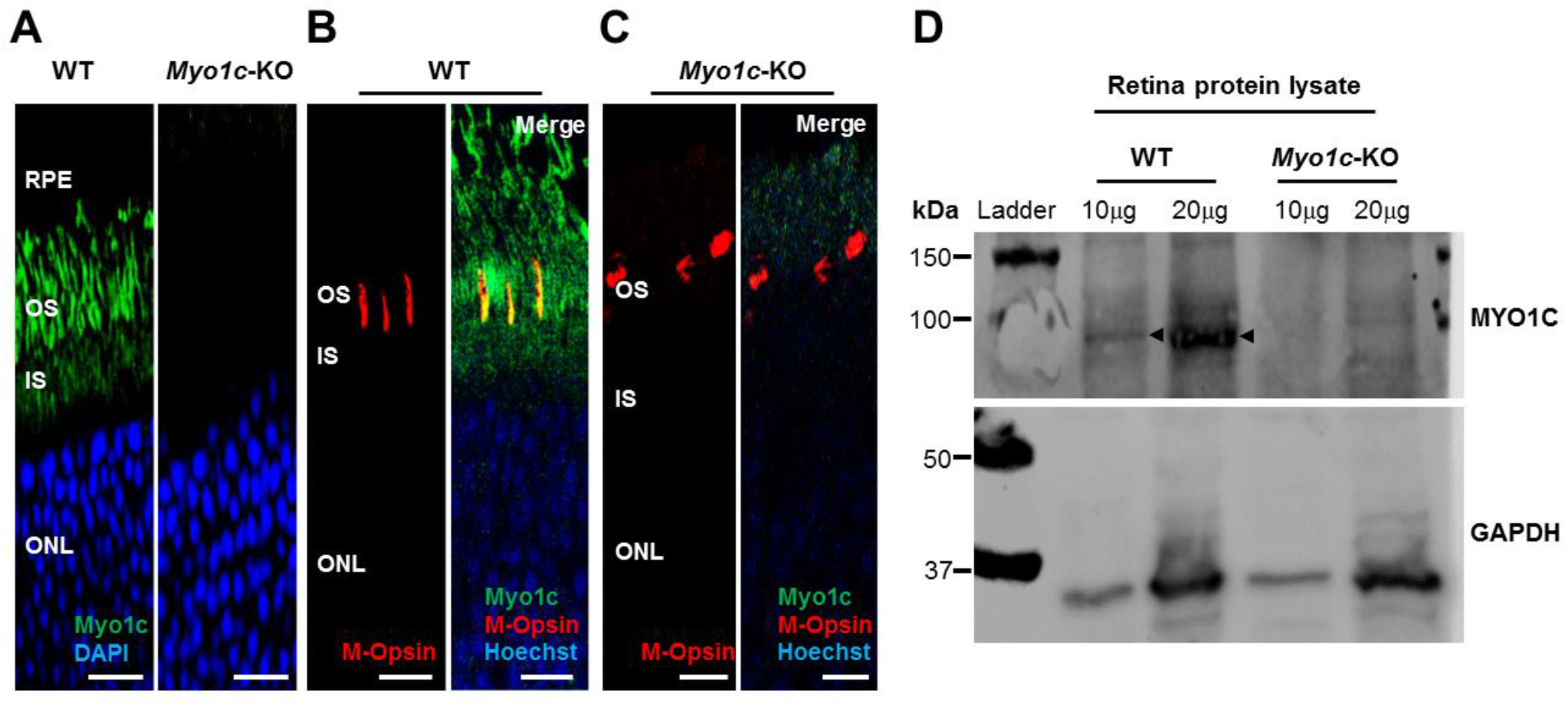
MYO1C localizes to photoreceptors in mouse retina: Eyes from adult wild-type (WT) and *Myo1c*-KO mice (*n*=8 mice per genotype; 50:50 ratio of male and female) were harvested and retina sections (*n*=5-7 sections per eye) were immunostained with an anti-MYO1C antibody (**a-c**), M-opsin antibody (**b, c**), followed by secondary (Alexa 488 or Alexa 594) antibody staining. MYO1C (green fluorescence), M-Opsin (red fluorescence), and DAPI or Hoechst (blue fluorescence). Figures in **a-c** are representative of retinal sections (*n*=5-7 sections per eye) imaged from *n*=8 animals per genotype. (**b, c**) Merge (orange) represents co-localization of MYO1C-488 (green) with M-Opsin-594 (red). RPE, retinal pigmented epithelium; OS, outer segments; IS, inner segments; ONL, outer nuclear layer. (**a-c**) Scale bar=50 µm. (**d**) Total protein isolated from WT (*n*=4) and *Myo1c*-KO (*n*=4) mouse retinas were pooled respectively and subjected to SDS-PAGE. Two different concentrations of protein (10μg and 20μg) were used. Blots were then probed with anti-Myo1c and Gapdh antibodies. Western blot analysis were repeated thrice. Arrows indicate MYO1C protein band in retinal lysates of WT mice.

**Fig. 2:**
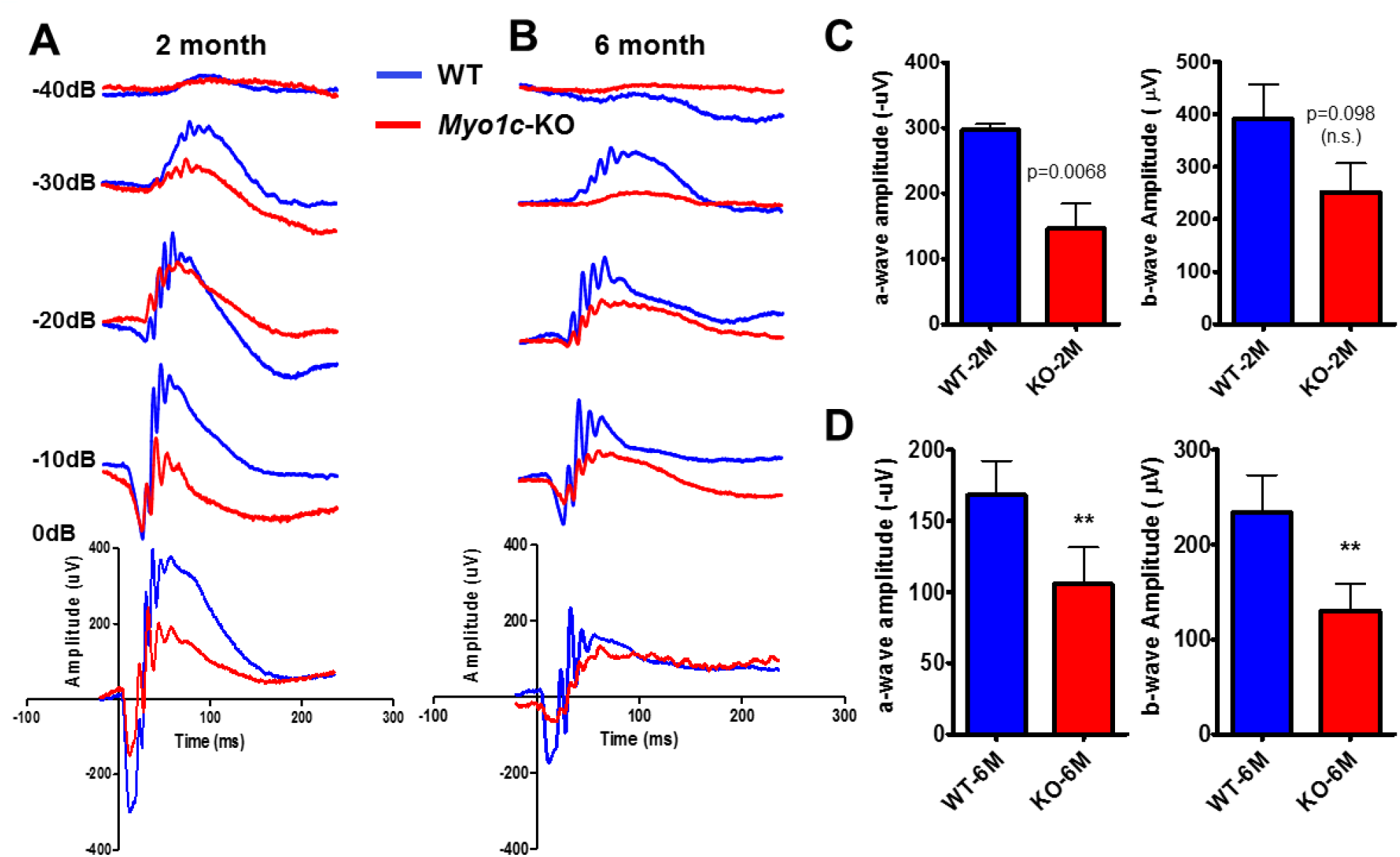
Genetic deletion of *Myo1c* in mice results in decreased visual function: Dark-adapted scotopic ERGs were recorded in response to increasing light intensities in cohorts of control wild-type/WT (blue bars, blue-traces) and *Myo1c*-KO (red bars, red-traces) mice of two month old (**a, c**), and six month old (**b, d**). Two-month-old *Myo1c*-KO mice had lower dark-adapted *a*- and *b*-wave amplitudes compared with controls (post-hoc ANOVA: *a*-waves, ^*^*p*<0.0068; *b*-waves, *p*<0.0098, n.s. not significant.), in particular at higher light intensities (−40, −30, −20, −10, 0 dB). Six-month-old *Myo1c*-knockout mice had lower dark-adapted *a*- and *b*-wave amplitudes compared with controls (post-hoc ANOVA: *a*-waves, ^**^*p*<0.005; *b*-waves, ^**^*p*<0.005), in particular at higher light intensities (−40, −30, −20, −10, 0 dB). Photoreceptor cell responses (*a*-waves), which drive the *b*-waves, were equally affected in 6-months old *Myo1c*-KO animals (both reduced on average between 38-45% of WT animals). Data are expressed as mean ± S.E. (*Myo1c*-KO mice and WT mice, *n*=8 per genotype and age-group; 50:50 ratio of male and female).

### Trafficking of Rod and Cone Visual Pigments in *Myo1c-*KO Mice

Since the phototransduction protein rhodopsin constitutes 85-90% of photoreceptor OS protein content [32], and as the ERG responses were impaired in *Myo1c*-KO mice, we hypothesized that the loss of MYO1C might have affected opsin trafficking to the photoreceptor OS. To test this hypothesis, we analyzed retinal sections from WT and *Myo1c*-KO mice (at 2 and 6 months of age; 5-7 retinal sections per eye from *n*=8 mice per genotype and age-group; 50:50 ratio of male and female), probing for rhodopsin, two types of cone opsins, medium wavelength R/G opsin (M-opsin) and short wavelength S-opsin, rod-specific phosphodiesterase 6b (Pde6b), rod-specific CNGA1, rod arrestin (ARR1), rod transducin (G-protein), and the general cone marker, PNA lectin. In WT mice at 2 and 6 months of age, rhodopsin localized exclusively to the rod OS (**Fig. 3a**). While majority of rhodopsin trafficked to the OS in two month old *Myo1c-*KO mouse retinas, some mislocalization to the base of the rod IS and the cell bodies in the outer nuclear layer (ONL) was noted (**Fig. 3a;** white arrows; rhodopsin levels within individual retinal layers were quantified and shown in **Figs. S3a-c**). This suggested incomplete opsin transport/trafficking to photoreceptor OS in the absence of MYO1C. An even more severe mislocalization of rhodopsin to the rod IS and within the ONL was observed in the 6-month old *Myo1c*-KO mice, suggesting a progressive retinal phenotype in the absence of MYO1C (**Fig. 3a;** rhodopsin expression within individual retinal layers were quantified and shown in **Figs. S3d-e**). Staining for the two cone opsins showed that the cone OS were shorter and mis-shaped by two months and this abnormality increased by six months of age (**Figs. 3b** and **3c**). Retinas stained for PNA lectin, showed progressively shorter and misshaped cone OS, indicating that cone OS structure was compromised in the absence of MYO1C as these mice aged (**Fig. 3d**). Cone visual arrestin in WT mice retina typically outlines the entire cell, OS, IS, cell body, axon, and cone pedicle. Staining for cone arrestin in *Myo1c*-KO animals (2 month of age) confirmed the short and mis-shaped appearance of the cone OS compared to WT retinas at similar ages (**Fig. 4a**, white arrows). In contrast, staining for Pde6b, a lipidated rod specific protein that trafficks to the OS independently of rhodopsin [33], showed normal trafficking and localization to the rod OS in both WT and *Myo1c*-KO retinas, at 2 month of age (**Fig. 4b**).

**Fig. 3:**
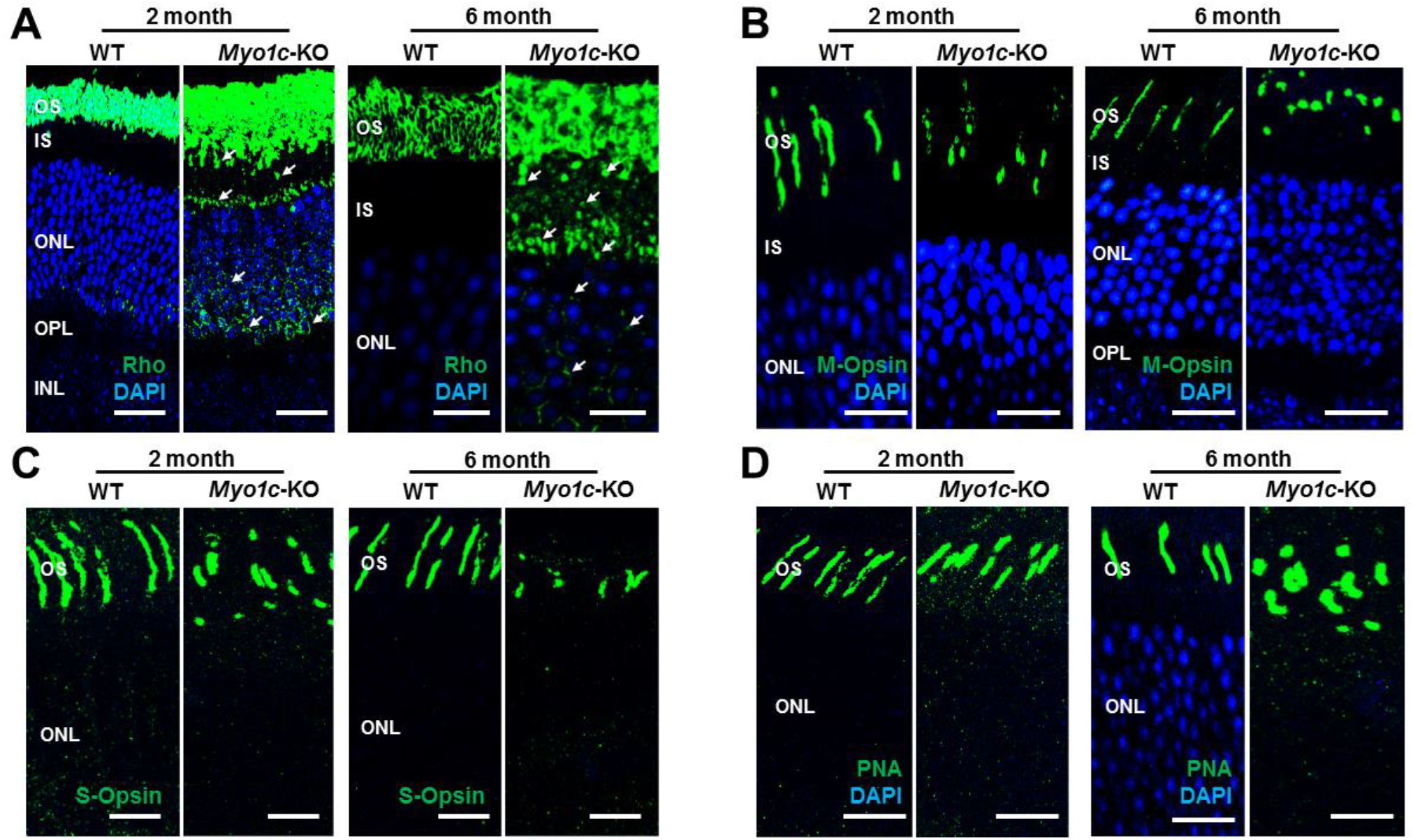
Immunohistochemical analysis of wild-type/WT and *Myo1c*-knockout mice retinas shows opsin trafficking defects: (**a**) Levels and localization of rhodopsin (Rho); **b**, red/green medium wavelength cone opsin (M-opsin); **c**, short wavelength cone opsin (S-opsin); **d**, PNA-488, were analyzed in two and six-months old WT and *Myo1c-KO* mice retinas. *Arrows* in panel ***a*** highlight rhodopsin mislocalization to IS and cell bodies in *Myo1c*-knockout mouse retinas. Images in panels **a-d** are representative of immunostained retinal sections (*n*=5-7 sections per eye) imaged from *n*=8 animals per genotype and age-group (50:50 ratio of male and female). Scale bar=75 µm (**a**); Scale bar=50 µm (**b, c, d**). OS, outer segments; IS, inner segments; ONL, outer nuclear layer; INL, inner nuclear layer; OPL, outer plexiform layer.

**Fig. 4:**
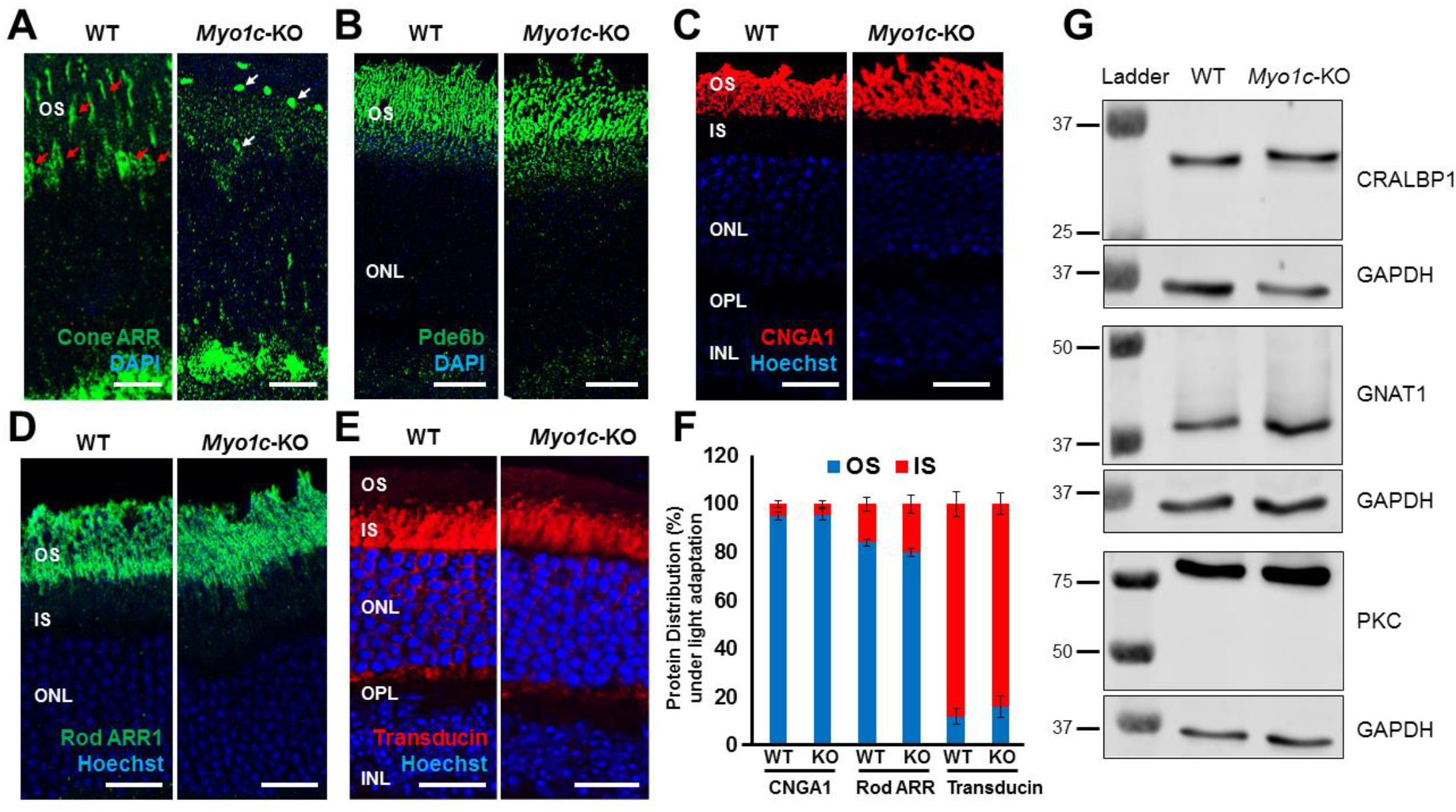
Immunohistochemical analysis of protein trafficking in photoreceptors of wild-type/WT and *Myo1c*-knockout mice retinas: Levels and localization of (**a**) cone arrestin (ARR), (**b**) Pde6b; (**c**) CNGA1; (**d**) Rod Arrestin (ARR1); and (**e**) G-protein (Transducin), were analyzed in WT and *Myo1c-KO* mice retinas to evaluate protein trafficking to photoreceptor OS. Red Arrows in **panel *a*** highlight cone photoreceptor nuclei and OS in WT mouse retinas that were significantly reduced or shorter respectively in *Myo1c-KO* animals (white arrows in **a**). Images in panels **a-e** are representative of immunostained retinal sections (*n*=5-7 sections per eye) imaged from *n*=8 animals per genotype and age group (50:50 ratio of male and female). **Panels a, b**, mice were 2-3 months of age. **Panels c-e**, mice were 3-4 months of age. (**f**) protein distribution (in %) of CNGA1, Rod ARR1, and Transducin, within the photoreceptor OS and IS, in light adapted mice. For quantification of protein distribution within retinal layers, 5-7 retinal sections from each eye (*n*=8 animals for each genotype) were analyzed using Image *J*. (g) Representative western blot (*n*=3 repeats) images of retinal proteins from 3-4 month old WT and *Myo1c-KO* mice (*n*=2 animals per genotype) showed no significant differences in protein expression of key retinal genes among genotypes. OS, outer segments; IS, inner segments; ONL, outer nuclear layer; INL, inner nuclear layer; OPL, outer plexiform layer; IPL, inner plexiform layer.

The CNG channels are also important mediators in the photoreceptor transduction pathways, and they require proper localization to the OS for normal photoreceptor cell function [5]. Additionally, the absence of CNGA1 or CNGB1 in mice led to decreased ERG responses and progressive rod and cone photoreceptor cell death [5]. Therefore, to rule out alternate mechanisms for the observed functional phenotypes in *Myo1c*-KO retinas, the retinas of WT and *Myo1c*-KO mice (3-4 months of age; 5-7 retinal sections per eye from *n*=8 mice per genotype; 50:50 ratio of male and female) were stained with the CNGA1 antibody. This analysis showed that even in the absence of MYO1C, both young and adult mice retinas showed no defects in the trafficking of CNGA1 protein to OS (**Fig. 4c;** CNGA1 protein distribution in photoreceptor layer quantified and shown in **Fig. 4f**).

The soluble proteins arrestin and transducin exhibit light-dependent trafficking, where in response to light, arrestin migrates to rod OS and transducin translocates to rod IS [34]. To test whether the loss of MYO1C affected rod arrestin (ARR1) and rod G-protein (transducin) localization, we performed IHC staining for these proteins in retinas of light adapted WT and *Myo1c-*KO mice (3-4 months of age; 5-7 retinal sections per eye from *n*=8 mice per genotype; 50:50 ratio of male and female). These analyses showed that in the presence of light, genetic loss of MYO1C had no negative effect on the trafficking of rod arrestin to the OS and G-protein to the IS and cell bodies in retinas of *Myo1c*-KO mice (**Figs. 4d** and **4e;** rod ARR1 and transducin protein distribution in photoreceptor layer quantified and shown in **Fig. 4f**). Using total protein lysates from retinas of WT and *Myo1c-*KO mice (3-4 months of age; four pooled retinas from *n*=2 mice per genotype) we analyzed protein expression of key retinal proteins in specific retinal cells: CRABLP1 (expressed in Müller cells), GNAT1 (expressed in photoreceptors), and PKCα (expressed in retinal bipolar cells). These analyses showed no significant differences in the expression of these genes in the inner or outer-retinal layers of *Myo1c*-KO mice when compared to WT mice, at 3-4 months of age (**Fig. 4g**). Although MYO1C could not be detected by immunohistochemical analysis in mouse RPE, functional MYO1C and *Myo1C* mRNA were reported in human RPE cells [35] and mouse RPE [36], respectively. Since elimination of the motor protein *Myo7a* in mouse leads to alterations in protein localization in the RPE (RPE65) [37], we stained retinas of young and adult WT and *Myo1c*-KO mice (5-7 retinal sections per eye from *n*=8 mice per genotype) with an anti-STRA6 antibody, another RPE-specific protein. This analysis showed that STRA6 expression and localization in the RPE was not affected in the absence of MYO1C (**Fig. S4**). Since MYO1C is known primarily as a motor protein with a protein trafficking function [14, 23], we next tested the hypothesis that its absence in photoreceptors of *Myo1c*-KO animals may contribute specifically to the loss of opsin trafficking to the photoreceptor OS.

### Native Cre+ mice showed no retinal phenotypes

To rule out any Cre+-mediated effects on retinal phenotypes observed in the *Myo1c*-KO;Cre+ animals, the eyes from native Cre+ mice (3-4 months old; *n*=3 animals) were harvested and subjected to similar histological and immunofluorescence analysis. As compared to age-matched WT mice retinas (*n*=3 animals), the retinas of Cre+ mice showed no retinal pathology or mislocalization of opsins (**Figs. S5a** vs. **S5b**). These analyses support the view that genetic loss of MYO1C affects key components of phototransduction specifically, and this is further manifested in defects in visual function.

### *Myo1c*-KO Mice demonstrated Photoreceptor OS Loss

To evaluate further if opsin mistrafficking is associated with structural changes to the retina, histological and transmission electron microscopy (TEM) analyses of retinal sections of young and adult WT and *Myo1c*-KO mice were performed. In histological sections of retinas (5-7 retinal sections per eye from *n*=8 mice per genotype and age), progressive shortening of rod photoreceptor OS was observed. The OS of adult *Myo1c*-KO mice at 6 months of age were shorter than the OS of *Myo1c*-KO mice at 2 months of age, which in turn were shorter than those in WT mice at similar ages (**Figs. 5a** and **5b;** OS lengths quantified from H&E sections and represented using spider-plots in **Figs. 5c** and **5d;** ^**\*\***^*p*<0.05). In comparison to WT mice, the photoreceptors in *Myo1c*-KO mice were less organized, especially in the 6-month old mice (**Fig. 5b**), suggesting that loss of MYO1C may progressively affect photoreceptor homeostasis. The retina outer nuclear layer (ONL) thickness between genotypes at both ages revealed no significant reduction in nuclear layers in *Myo1c*-*KO* animals compared to WT mice (ONL thickness quantified from H&E stained sections and represented using spider-plots in **Figs. 5e** and **5f**).

**Fig. 5:**
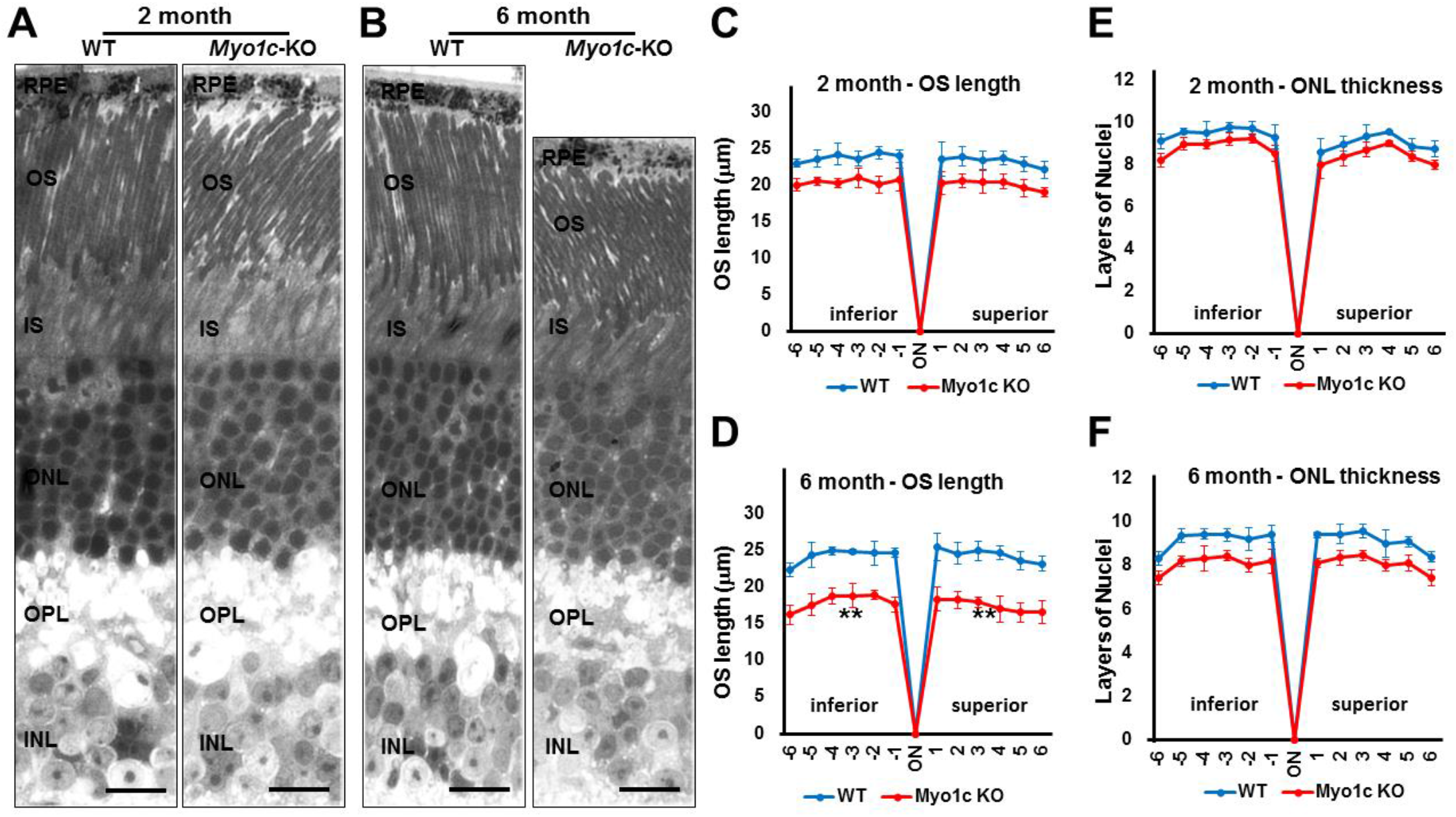
Histological analysis shows reduced photoreceptor OS lengths in *Myo1c*-KO mice retinas: (**a, b**) Retinas from 2 and 6 month old WT and *Myo1c-KO* mice were sectioned using an ultra-microtome and semi-thin plastic sections were obtained to evaluate pathological consequences of MYO1C loss. Quantification of OS lengths from H&E sections (**c**, two month old mice; **d**, 6 month old mice) and ONL thickness (**e**, two month old mice; **f**, six months old mice) using “spider graph” morphometry. The OS lengths and total number of layers of nuclei in the ONL of from H&E sections through the optic nerve (ON; 0 μm distance from Optic Nerve and starting point) was measured at 12 locations around the retina, six each in the superior and inferior hemispheres, each equally at 150μm distances. RPE, retinal pigmented epithelium; OS, outer segments; IS, inner segments; ONL, outer nuclear layer; INL, inner nuclear layer; OPL, outer plexiform layer; IPL, inner plexiform layer; GCL, ganglion cell layer. Retinal sections (*n*=5-7 sections per eye) from *n*=8 mice each genotype and time-point (50:50 ratio of male and female) were analyzed. Two-way ANOVA with Bonferroni posttests compared *Myo1c*-KO mice with WT in all segments. \*\**p*<0.005, for OS length in only 6 month old *Myo1c*-KO mice compared to WT mice; and n.s. (not significant) for ONL thickness in both 2 month and 6 month old *Myo1c*-KO animals, compared to WT mice). (a, b) Scale bar=100μm.

### Ultrastructural TEM analysis showed shorter photoreceptor OS in *Myo1c*-KO mice

To evaluate the structure of rod photoreceptors, ultrastructural analysis using TEM was performed (*n*=6 retinal sections per eye from *n*=8 mice per genotype and age). While the rod photoreceptor OS in the WT mice showed normal elongated morphology, they appeared slightly shorter in *Myo1c*-KO mice at two months of age (^*^*p*<0.05; **Fig. 6a;** rod OS lengths quantified in **Fig. 6e**). Specifically, comparing *Myo1c-*KO with WT mouse rod OS lengths at six months of age demonstrated that OS segment lengths in *Myo1c* retinas were significantly (36-45%) shorter than those of WT mice (^**^*p*<0.005; **Fig. 6b;** rod OS lengths quantified in **Fig. 6e**). Ultrastructurally, the cone OS in the *Myo1c*-KO mouse retina were shorter and had lost their typical cone shape (**Fig. 6c** vs. **6d;** cone OS lengths quantified in **Fig. 6f**), confirming the mis-shaped cone OS phenotype identified by immunohistochemistry (**Figs. 3b-d**). These results suggest that the lack of MYO1C resulted in progressively severe opsin mislocalization (**Figs. 3a-d**), and shorter photoreceptor OS (**Figs. 5** and **6**), thus supporting the observed decrease in visual function by ERG (**Fig. 2**).

**Fig. 6:**
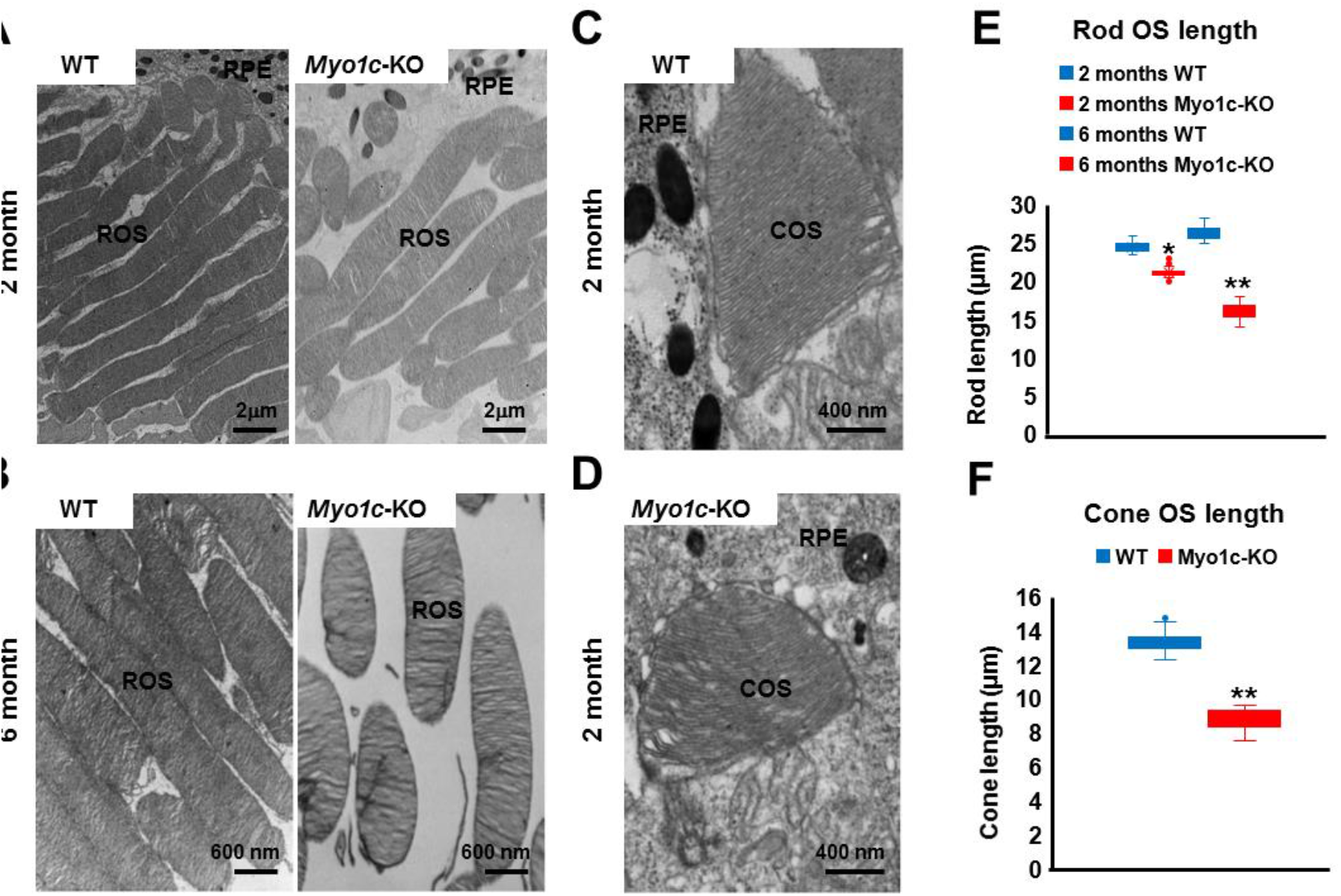
Ultrastructural analysis of rods and cone photoreceptors using transmission electron microscopy (TEM): Representative TEM images of rod photoreceptors from two month (**a**) and six month (**b**) old WT and *Myo1c*-*KO* mice, are presented. Representative images of cone photoreceptors from 2 month old WT (**c**) and *Myo1c*-*KO* (**d**) mice. (**a**) Scale bar=2μm (**b**) Scale bar=600 nm (**c, d**) Scale bar=400 nm. Data is representative of *n*=6 retinal sections per eye from *n*=8 mice per genotype and time-point. (**e**) Rod OS (ROS) length in WT animals were measured and compared to *Myo1c*-KO animals. (**f**) Cone OS (COS) length in WT animals were measured and compared to *Myo1c*-KO animals.^*^*p*<0.05; ^**^*p*<0.005. RPE, retinal pigmented epithelium.

### Molecular inhibition of MYO1C motor function by PCIP in mice affected Opsin trafficking

To confirm a direct role for the motor protein MYO1C in opsin trafficking and that the loss of MYO1C contributed specifically to opsin mistrafficking in photoreceptors, we molecularly inhibited MYO1C in vivo using PClP (pentachloropseudilin) which specifically can inhibit the motor activity of MYO1C [38-41]. To achieve this, a single dose of PCIP in DMSO (5mg/kg) was injected retro-orbitally into the right eye of WT animals (*n*=2). A control set of WT animals (*n*=2) received vehicle control/diluent (DMSO) under similar conditions. Post 7-8 hours injection, mice were euthanized and both eyes were harvested and fixed in PBS buffered 4% paraformaldehyde. Retinal cell phenotype and opsin trafficking in PCIP and DMSO injected animals were assessed using retinal histology and immunofluorescence to assess rod and cone opsin trafficking to the OS. In photoreceptors of mice eyes injected with the vehicle control DMSO, rhodopsin localized exclusively to the rod OS (**Fig. 7a**). In contrast, the mice injected with PCIP showed mislocalization of rhodopsin to the base of the rod IS and the cell bodies in the ONL, demonstrating incomplete opsin trafficking to the photoreceptor OS in PCIP injected mice retinas (**Fig. 7a;** white arrows). Observation of retinal phenotypes in the left eyes of these animals indicated that injected PCIP was systemically distributed (**Fig. 7a)**. Staining for the R/G cone opsins (M-opsins) in retinas of these animals showed that in comparison to the control mice the cone OS of PCIP injected mice were shorter and lost their typical elongated cone morphology (**Fig. 7b**). The histological analysis of retinal sections showed significant shortening of photoreceptor OS in mice retinas injected with PCIP (**Fig. 7c**). Additionally, the quantification of retinal ONL thickness in PCIP injected mice showed that ONL was slightly thinner in comparison to control mice (ONL thickness quantified from H&E sections and represented using spider-plots in **Fig. 7d)**. OS lengths were significantly shorter in PCIP injected mice (^*^p<0.05; **Fig. 7c;** quantified from H&E sections and represented using spider-plots in **Fig. 7e**). These retinal phenotypes were similar to those observed in the retinas of *Myo1c*-KO mice at two and six months (**Figs. 3, 5a**, and **6**). Collectively, these results indicate that MYO1C is critical for opsin trafficking to photoreceptor OS and its loss specifically affects opsin trafficking, photoreceptor cell homeostasis, and visual function.

**Fig. 7:**
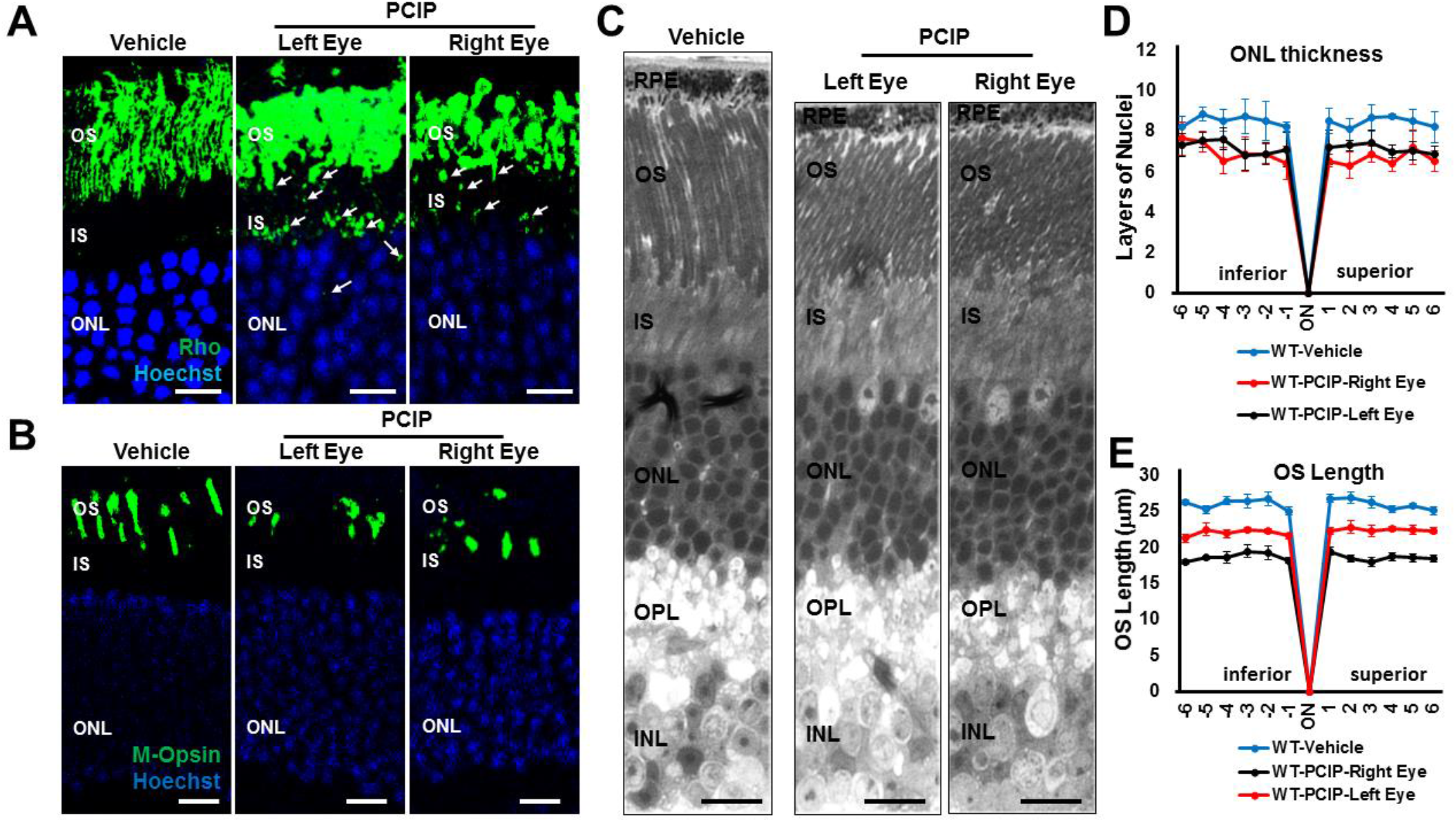
Histological and immunohistochemical analysis of mice injected with PCIP: The *Myo1c* specific inhibitor PCIP (pentachloropseudilin) or vehicle control DMSO was injected retro-orbitally into the right eye of WT animals (*n*=2 per treatment). (**a**) Levels and localization of rhodopsin (Rho) and (**b**), red/green opsins (M-opsin). (**c**) Semi-thin plastic sections of retina from mice were obtained to evaluate pathological consequences of PCIP treatment. (**d**) Quantification of ONL thickness and (**e**) photoreceptor OS lengths using “spider graph” morphometry, from H&E sections. The OS length and total number of layers of nuclei in the ONL of semi-thin plastic sections through the optic nerve (ON; 0μm distance and the starting point) was measure at 12 locations around the retina, six each in the superior and inferior hemispheres, each equally at 150μm distance. Retinal sections (*n*=10-12 sections per eye) from *n*=2 mice of each treatment group were evaluated for ONL thickness and OS lengths. (**a, b**) Scale bar=50µm (**c**) Scale bar=100 µm. RPE, retinal pigmented epithelium; OS, outer segments; IS, inner segments; ONL, outer nuclear layer; INL, inner nuclear layer; OPL, outer plexiform layer.

### MYO1C Directly Interacted with Rhodopsin

Since the loss of MYO1C resulted in retinal function defects with significant alterations in the localization of opsins, we next evaluated whether MYO1C exerted this effect through a physical interaction with rhodopsin. Immunoprecipitation analysis using WT and *Myo1c*-KO mice retinas (*n*=6 retinas pooled from *n*=3 animals per genotype, respectively) demonstrated that rhodopsin was pulled down using MYO1C antibody, and this interaction was confirmed in a reciprocal fashion (**Fig. 8a;** Co-IP flow-chart schematic shown in **Fig. S6**). Using a baculovirus-produced purified recombinant mouse MYO1C protein in an overlay assay, we demonstrated that MYO1C directly interacted with rhodopsin, where opsin was immunoprecipitated both from mouse retinal lysate or HEK293 cells transfected with pCDNA3 Rod Opsin in order to overexpress Rhodopsin (schematic representation in **Fig. S6**). Immunoprecipitated rhodopsin was subjected to western blotting and probed with purified recombinant full-length MYO1C (MYO1C FL; **Fig. 8b** and schematic in **Fig. S6**) or GFP-MYO1C-790-1028 (MYO1C tail domain, also known as the cargo domain; **Fig. 8c** and **Fig. S6**) [13]. Post-incubation, the interaction of immobilized rhodopsin to Myo1c was probed using a MYO1C antibody. The immunoblot analysis of the over-layered MYO1C showed significant binding of both MYO1C proteins, full-length and the tail domain, at the rhodopsin band, indicating a direct interaction between the two proteins (**Figs. 8b** and **8c**). Interestingly, the interaction of MYO1C was noted with various multimers of rhodopsin, which further indicated that opsin is a cargo for MYO1C (arrows in **Figs. 8b** and **8c**).

**Fig. 8:**
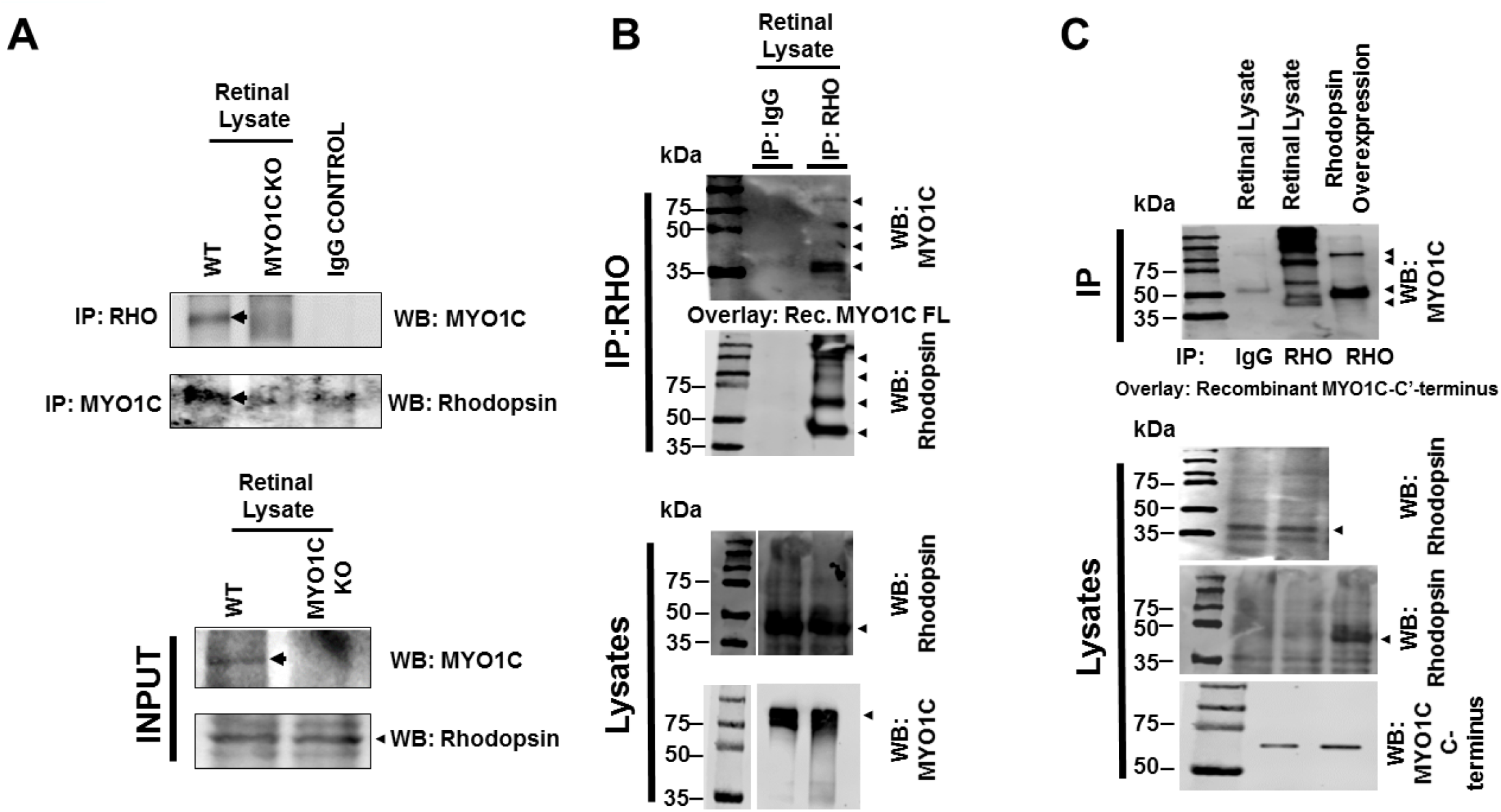
Rhodopsin is a direct cargo for MYO1C: (**a**) Mice retinal protein lysates were isolated from *Myo1c*-KO and wild type (WT) mice (6 retinas pooled from *n*=3 mice per genotype) and subjected to co-immunopreciptation analysis. Rhodopsin was co-immunoprecipitated with MYO1C antibody (top panel). In a reciprocal manner, MYO1C was co-immunoprecipated with Rhodopsin antibody (bottom panel). (**b**) Using Rhodopsin antibody, Rhodopsin (RHO) was immunoprecipitated either from mice retinal lysates or from HEK293 cells (transfected with pCDNA rhodopsin plasmid) where Rhodopsin was overexpressed. The immunoprecipated Rhodopsin was separated using SDS-PAGE and transferred to nitrocellulose membranes. The rhodopsin bound to nitrocellulose membrane was then incubated with 5ug of purified recombinant active MYO1C-full length or MYO1C C-terminal cargo domain protein (**c**) generated from a baculovirus expression system. To analyze if MYO1C binds to immobilized Rhodopsin, blots were washed and western blotted with MYO1C antibody. A positive signal with MYO1C showed direct binding of MYO1C to various Rhodopsin multimers (arrows) present in the retinal lysate and overexpressed pCDNA3-Rhodopsin in HEK293 cells.

### Genetic Deletion of *Myo1c* did not affect Systemic Organs in Mice

Finally, to determine if the global deletion of *Myo1c* affected other organs, we harvested major systemic organs, including liver, heart, and kidney of 2 month old *Myo1c*-KO and WT mice (*n*=4 per genotype), and performed histological analyses. Notably, *Myo1c-*KO mice developed and reproduced normally with no observable histological differences between the control and *Myo1c*-KO genotypes (**Figs. S7a-c**). To further confirm that there were no functional defects in these systemic organs, we performed ECHOcardiogram (heart function), quantified protein/albumin levels in urine (kidney function), and measured Alanine Aminotransferase/ALT enzyme levels (liver function), in *Myo1c*-KO mice (*n*=4 mice per individual functional analysis) and compared these values to their WT littermates (*n*=4 mice per individual functional analysis). All of these analyses showed no pathological defects in systemic organs of *Myo1c*-KO animals when compared to the age-matched WT littermates (**Figs. S7a’-c’** and **S8**). Overall, these results indicate that except for the retinal phenotypes, *Myo1c*-KO animals retained normal physiology of the systemic organs examined.

## Discussion

The trafficking of the G-protein coupled receptor (GPCR) Type II Opsins from the photoreceptor IS to the OS represents a critical event in the initiation of phototransduction for visual function in vertebrates. Our work identified for the first time an unconventional motor protein, MYO1C, as a novel trafficking regulator of both rod and cone opsins to the photoreceptor OS in mice. In this study, based on MYO1C localization within the IS and OS of photoreceptors, and using a whole-body *Myo1c*-KO mouse model, we functionally identified MYO1C as a novel component of retinal physiology and was specifically found to be involved in photoreceptor cell function. Retinal analysis of *Myo1c*-KO mice identified opsins as novel cargo for MYO1C. In the absence of MYO1C, both young and adult *Myo1c*-KO mice showed impaired opsin trafficking, where rhodopsin was retained in the photoreceptor IS and the cell bodies. In contrast, cone opsins showed no retention in the cell body or mistrafficking to other retinal cell layers, although staining patterns revealed deformed cone OS shapes. These two phenotypes manifested as a progressive decline of visual responses in the rod ERGs and shorter photoreceptor OS lengths as *Myo1c*-KO animals aged, indicating a progressive retinal phenotype. Interestingly, trafficking of other OS proteins (CNGA1, arrestin, and transducin) were largely unaffected in the absence of MYO1C. The genetic deletion of *Myo1c* only affected retina, and the other systemic organs examined, including heart, liver, and kidney, remained unaffected. Use of PClP as an allosteric inhibitor of MYO1C ATPase and motor activity resulted in retinal phenotypes similar to those observed in *Myo1c*-KO mice and thus confirmed that MYO1C plays a critical role in the trafficking of opsin to the photoreceptor OS. Overall, our data points to a novel mechanism by which MYO1C regulates opsin trafficking from the photoreceptor IS to OS, a critical event for photoreceptor function and long-term photoreceptor cell homeostasis. Our study identifies an unconventional motor protein MYO1C as an essential component of mammalian photoreceptors, where it plays a canonical role in promoting opsin trafficking and maintaining normal visual function.

### MYO1C and Other Opsin Trafficking Proteins

*Myo1c*-KO mice exhibited rhodopsin mislocalization similar to that of *Rpgr*^-/-^, *Myo7a*^Sh1^, *Rp1*^-/-^, *Kinesin II*^-/-^, and *Tulp1*^-/-^ mutant mice [1-9]. Since MYO1C primarily localized to photoreceptor IS and OS, is known to be involved in protein trafficking, and uses actin as a track [14, 23], we hypothesized that MYO1C participates in the movement of opsins from IS to the OS of photoreceptors. This hypothesis was supported by the observation that the rod opsins were mislocalized to IS and cell bodies. Defective assembly of cone OS in *Myo1c-*KO mice suggests that this phenotype is caused by an aberrant protein transport with OS degeneration as a secondary event. The normal ultrastructure of photoreceptors in our *Myo1c*-KO mice suggests that the retinal abnormalities in these animals were not due to structural defects in photoreceptors per se, but instead were induced by aberrant motor function leading to opsin mislocalization.

### MYO1C Contributed to Phototransduction and Retinal Homeostasis

The opsin molecules and other phototransduction proteins are synthesized in the cell body of the photoreceptor [42, 43]. They are then transported to the distal IS [44] and subsequently to the OS. Little is known about these transport processes and the molecular components involved in this process [1-9]. The localization of MYO1C in the rod photoreceptors’ IS and OS, and in cone OS, suggested that opsins may utilize this molecular motor for transport to the OS. The immunohistochemical analysis of *Myo1c*-KO animals indicated that while rod and cone opsins trafficked to the OS, significant mislocalization was noted for rhodopsin in the IS and cell bodies in the ONL (**Fig. 2**). Since they represent plasma membrane structural proteins, cone opsins presumably contribute to the cone OS stability and rhodopsin to the rod OS formation and stability [7]. Hence, photoreceptor OS shortening/degeneration in *Myo1c-*KO mice may be attributed, in large part, to the mistrafficking of opsins to the IS or a progressive reduction of opsins in the OS membrane. Notably, the pattern of opsin mislocalization observed in *Myo1c*-KO mice closely resembled the retinal phenotype observed in our previously reported *Tulp1*-KO mice [4, 45], *Cnga3*^-/-^ mice [5], *Lrat*^-/-^ and *Rpe65*^-/-^ mice [3, 8, 9], GC1-KO mice [1, 6], and, to some extent in CFH (complement factor H)-KO animals [2]. Importantly, in all these studies, photoreceptor OS were unstable, and significant degeneration was noted. However, because 85-90% of OS protein is rhodopsin, the mislocalization of other less abundant proteins cannot be ruled out in the photoreceptors of *Myo1c*-KO mice.

### Contributions from Other Motor proteins in Opsin trafficking

Although this study demonstrates mistrafficking of opsins due to a loss of MYO1C, the majority of opsin was still correctly localized, suggesting that contribution or compensation from other myosins cannot be ruled out. Nevertheless, the contributions from MYO1C were highly significant as its genetic deletion showed specific physiological defects in mouse retinas. It is likely that some redundancy exists among molecular motors, and several known candidates might compensate for the lack of MYO1C in photoreceptor function. However, the Qpcr analysis of the retinas from WT and *Myo1c-KO* mice did not suggest compensation from other family myosin 1 members (**Fig. S9**). Interestingly, the upregulation of Myo1f in our study was unable to rescue the Myo1c retinal phenotype suggesting that Myo1f is unable to compensate for the functional loss of Myo1c in retina (**Fig. S9**). However, compensation by other motor proteins, including the members of kinesin superfamily [46, 47], myosin VIIa, and conventional myosin (myosin II) [48, 49], which have also been detected in the RPE and retina, cannot be ruled out and need further investigation. Overall, these results support a direct role for MYO1C in opsin trafficking in the photoreceptor cells of the retina and provide evidence that defective protein transport pathways are a pathologic mechanism responsible for OS degeneration and decreased visual function in these mice.

## Methods

### Materials

All chemicals, unless stated otherwise, were purchased from Sigma-Aldrich (St. Louis, MO, USA) and were of molecular or cell culture grade quality.

### *Myo1c*-knockout (*Myo1c*-KO) Mouse Model

Mice were kept with *ad libitum* access to food and water at 24°C in a 12:12 h light–dark cycle. All mice experiments were approved by the Institutional Animal Care and Use Committee (IACUC protocol #00780; G.P.L.) of the Medical University of South Carolina, and performed in compliance with ARVO Statement for the use of Animals in Ophthalmic and Vision Research. We have previously generated *Myo1c* transgenic mice (*Myo1cfl/fl*) in C57BL/6N-derived embryonic stem cells, flanking exons 5 to 13 of the mouse *Myo1c* gene, which has allowed us to specifically delete all *Myo1c* isoforms in a cell-specific manner [29]. Here a complete *Myo1c*-knockout was generated by crossing *Myo1cfl/fl* mice with an F-actin Cre mouse strain (B6N.FVB-Tmem163Tg(ACTB-cre)2Mrt/CjDswJ) obtained from Jackson Labs. We will refer to the *Myo1cfl/fl* x f-actin Cre cross as *Myo1c* knockout (*Myo1c*-KO) mice. For this study, the *Myo1c*-KO mice were crossed onto a C57BL/6J background to avoid potential problems with the *Rd8* mutation (found in C57BL/6N lines) [50]. Equal numbers of male and female mice (50:50 ratio) were used per group and time-point.

### Immunohistochemistry and Fluorescence Imaging

Light-adapted mice were euthanized and eyes immediately enucleated. Eyes were fixed in 4% paraformaldehyde buffered with 1X PBS for 2 hours at 4°C using established protocols [58]. After fixation, samples were washed in 1X PBS and embedded in paraffin and processed (MUSC Histology core facility). Sections (10 µm) were cut and transferred onto frost-free slides. Slide edges were lined with a hydrophobic marker (PAP pen) and deparaffinized using xylene and processed through ethanol washes before blocking for 1-2 hours at RT. Blocking solution (1% BSA, 5% normal goat serum, 0.2% Triton-X-100, 0.1% Tween-20 in 1X PBS) was applied for 2 hours in a humidified chamber. Primary antibodies were diluted in blocking solution as follows: anti-rhodopsin (1:500, Abcam, 1D4), anti-Myo1c (1:100), cone-arrestin (1:250, Millipore-Sigma, St. Louis, MO), conjugated PNA-488 (1:2000, Molecular Probes, Eugene, OR), anti-red/green cone opsin (M-opsin; 1:500; Millipore, St. Louis, MO), anti S-opsin (1:500, Millipore/Sigma, St. Louis, MO), ZO1 (1:2000, Invitrogen), Pde6b (1:300, ThermoFisher), CNGA1 (1:250, Abcam), rod arrestin (1:250, Invitrogen), Stra6 (1:250, Millipore-Sigma), CRALBP (1:100, Invitrogen), rod transducin (1:250, Santa Cruz), and 4′,6-diamidino-2-phenylendole (DAPI; 1:5000, Invitrogen) or Hoechst (1:10,000, Invitrogen) was used to label nuclei. All secondary antibodies (Alexa 488 or Alexa 594) were used at 1:5000 concentrations (Molecular Probes, Eugene, OR). Optical sections were obtained with a Leica SP8 confocal microscope (Leica, Germany) and processed with the Leica Viewer software. All fluorescently labeled retinal sections on slides were analyzed by the BioQuant NOVA Prime Software (R & M Biometrics, Nashville) and fluorescence within individual retinal layers quantified using Image *J* or Fiji (NIH).

### Measurement of Photoreceptor ONL thickness and OS lengths

The lengths of the photoreceptor OS in WT and *Myo1c*-KO animals (from H&E sections of retinas) were imaged (Keyence BZ-X800 microscope) and measured at 12 consecutive points (at 150 μm distances) from the optic nerve (ON). The OS length was measured from the base of the OS to the inner side of the retinal pigment epithelium. The total number of layers of nuclei in the ONL of retinal sections through the optic nerve (ON) was imaged (Keyence BZ-X800 microscope) and measured at 12 locations around the retina, six each in the superior and inferior hemispheres, starting at 150 μm from the ON. Retinal sections (*n*= 5-7 retinal sections per eye) from *n*=8 mice for each genotype and time-point were analyzed. Two-way ANOVA with Bonferroni post-tests compared *Myo1c*-KO to WT mice, at each segment measured.

### PCIP (Pentachloropseudilin) retro-orbital injections

We and others have previously shown that the natural compound pentachloropseudilin (PClP) acts as an allosteric inhibitor of MYO1C ATPase and motor activity [38-41, 51-53]. To test whether the inhibition of MYO1C function by PCIP affects opsin trafficking, PCIP was retro-orbitaly injected (5mg/kg body weight) into the right eye of two-month old WT animals (*n*=2). At this concentration, PCIP was observed to inhibit all *Myo1c* isoforms without affecting non-*Myo1* Myosins [35, 38, 40, 41, 51-56]. The control set of WT animals (*n*=2) received retro-orbital injections of vehicle control (DMSO). Post 7-8 hours injection, mice were euthanized, both eyes were harvested, and fixed in 4% PFA for histological analysis.

### ERG Analysis

Dark-adapted WT and *Myo1c*-KO mice (50:50 ratio of male and female; *n*=8 each genotype) at 2 month of age (young mice; early time-point), and 6 month of age (end time-point) were anesthetized by intraperitoneal injection of a ketamine/xylene anesthetic cocktail (100 mg/kg and 20 mg/kg, respectively) and their pupils were dilated with 1% tropicamide and 2.5% phenylephrine HCl. ERGs were performed under dim red-light in the ERG rooms in morning (8am-12noon). Scotopic ERGs were recorded with a computerized system (UTASE-3000; LKC Technologies, Inc., Gaithersburg, MD, USA), as previously described [57-59].

### TEM analysis of retinas

Eyecups at the indicated time-points were harvested and fixed overnight at 4°C in a solution containing 2% paraformaldehyde/2.5% glutaraldehyde (buffered in 0.1M cacodylate buffer). Samples were rinsed in the buffer (0.1 M cacodylate buffer). Post-fixative 2% OsO_4_/0.2 M cacodylate buffer 1 hour at 4°C, followed by 0.1 Mcacodylate buffer wash. The samples were dehydrated through a graded ethanol series and then embedded in epon (EMbed 812; EM Sciences). For TEM analysis, each eye (*n*=6 individual eyes from *n*=6 animals of each genotype) was cut in half before embedding in epon blocks. Sections were parallel to the dorsoventral meridian and near the optic nerve (ON). The cured blocks were sectioned at 0.5 microns (semi-thin plastic sections) and stained with 1% toluidine blue to orient the blocks to the required specific cell types. The blocks were trimmed to the precise size needed for ultrathin sectioning. The blocks were cut at 70 nm and gathered on 1-micron grids. The grids were air-dried, stained with uranyl acetate for 15 minutes, lead citrate for 5 minutes, and rinsed between each stain. They were allowed to dry and imaged with a JEOL 1010. Images were acquired with a Hamamatsu camera and software. All samples were processed by the Electron Microscopy Resource Laboratory at the Medical University of South Carolina, as previously described [57].

### Western Blot Analysis and Densitometry

Total protein from cells or mouse tissues (*n*=3 per genotype) were extracted using the M-PER protein lysis buffer (ThermoScientific, Beverly, MA) containing protease inhibitors (Roche, Indianapolis, IN). Approximately 25 μg of total protein was electrophoresed on 4-12% SDS-PAGE gels and transferred to PVDF membranes. Membranes were probed with primary antibodies against anti-*Myo1c* (1:250), CRALBP (1:100, Invitrogen), Rod transducin (1:250, Santa Cruz), PKCα (1:500, Novus Biologicals), and β-Actin or Gapdh (1:10,000, Sigma) in antibody buffer (0.2% Triton X-100, 2% BSA, 1X PBS) [54,55,70]. HRP conjugated secondary antibodies (BioRad, Hercules, CA) were used at 1:10,000 dilution. Protein expression was detected using a LI-COR Odyessy system, and relative intensities of each band were quantified (densitometry) using Image *J* software version 1.49 and normalized to their respective loading controls. Each western blot analysis was repeated thrice.

### Co-immunoprecipitation (co-IP) Assays

Co-immunoprecipitation of endogenously expressed proteins (MYO1C and rhodopsin) was performed using mouse retinal extracts. Six retinas of each genotype (*n*=3 animals of WT and *Myo1c*-KO) were used for extraction of retinal proteins in 250 µL of RIPA buffer (phosphate-buffered saline [PBS] containing 0.1% sodium dodecyl sulfate [SDS], 1% Nonidet P-40, 0.5% sodium deoxycholate, and 100 mM potassium iodide) with EDTA-free proteinase inhibitor mixture (Roche Molecular Biochemicals). Lysates were cleared by centrifugation at 10000 rpm for 10 min at 4°C. The prepared lysates were further incubated with anti-Myo1c, anti-rhodopsin, and mouse/rabbit IgG overnight at 4°C and further with protein G-coupled agarose beads (ROCHE) for 1-2 h. Beads were then collected by centrifugation at 3000 rpm for 5 min at 4°C, extensively washed in 1X PBS, and resuspended in SDS gel loading buffer. The proteins were separated on a 10% SDS-PAGE, transferred to a PVDF membrane, and analyzed by immunoblotting with the corresponding antibodies (**Fig. S6**).

### Overlay direct binding assay

Rhodopsin protein was expressed in HEK293 cells using transient transfection (pcDNA3 rod opsin construct, a gift from Robert Lucas (Addgene plasmid # 109361, http://n2t.net/addgene:109361; RRID:Addgene_109361) [57] and immunoprecipitated from the cell lysates using an anti-rhodopsin antibody (Abcam). In parallel, rhodopsin was similarly immunoprecipitated from the mouse retinal lysate. The immunoprecipitated complexes were separated on SDS-PAGE gel and transferred to PVDF membrane. The membrane was then probed by overlaying it with 5 µg of baculovirus-produced and purified recombinant full-length MYO1C FL or GFP-MYO1C-790-1028 (tail domain, also known as the cargo domain [13]) protein, by incubating at 4°C for 4 h. Following incubation, the membranes were western blotted with MYO1C antibody to detect the direct binding of MYO1C to the rhodopsin bands. The location of rhodopsin on the membranes was marked by separately probing these membranes with an anti-rhodopsin (1:500, Millipore Sigma) antibody (**Fig. S6**).

### Quantitative Real Time-PCR

RNA was isolated from retinas of WT and *Myo1c*-KO animals using Trizol reagent, and processed as described previously [55,70]. One microgram of total RNA was reverse transcribed using the SuperScript II cDNA Synthesis Kit (Invitrogen, Eugene, OR). Quantitative Real-Time PCR (qRT-PCR) was carried out using SYBR green 1 chemistry (BioRad, Hercules, CA). Samples for qRT-PCR experiments were assayed in triplicate using the BioRad CFX96 Q-PCR machine. Each experiment was repeated twice (*n*=6 reactions for each gene), using newly synthesized cDNA.

### Liver function tests using Alanine Aminotransferase (ALT) assays

To extract total protein, liver tissue from WT or *Myo1c*-KO mice (pooled livers *n*=4 mice per genotype respectively) were homogenized in RIPA buffer on ice and then centrifuged at 14,000 rpm at 4°C for 10 min. Supernatant was collected, and the protein concentration was estimated using the Bio-Rad Protein Assay Dye Reagent (Sigma). 10 µl of liver lysate was transferred to 96-well plate and ALT was measured using a microplate-based ALT activity assay kit (Pointe Scientific, Cat. A7526). Five biological replicates were used in the assay.

### Heart function tests using Echocardiographic (ECHO) analyses

Echocardiographic (ECHO) analysis was performed on adult wildtype (WT) and *Myo1c*-KO animals (*n*=4 per genotype) at the MUSC Cardiology Core Facility. For ECHO experiments, mutant and wild-type littermate controls were anesthetized in an induction chamber with 5% isoflurane in 100% oxygen. They were removed and placed on a warming table where anesthesia was maintained via nose cone delivery of isoflurane (1% in 100% oxygen). They were placed in the supine position, and the thoracic area was shaved. The limbs were taped to the platform to restrict animal movement during echocardiography acquisition. This also provided a connection to ECG leads embedded in the platform. Sonography gel was applied to the chest and echocardiographic measurements of the peristernal long axis and short axis of the heart were acquired to derive the systolic and diastolic parameters of heart function. ECHO measurements were estimated using vevo 2100 instrumentation.

### Statistical Analysis

Data were expressed as means ± standard deviation by ANOVA in the Statistica 12 software (StatSoft Inc., Tulsa, Oklahoma, USA). Differences between means were assessed by Tukey’s honestly significant difference (HSD) test. *P*-values below 0.05 (*P*<0.05) were considered statistically significant. For western blot analysis, relative intensities of each band were quantified (densitometry) using the Image *J* software version 1.49 and normalized to the loading control β-actin. The qRT-PCR analysis was normalized to 18S RNA, and the ΔΔCt method was employed to calculate fold changes. Data of qRT-PCR were expressed as mean ± standard error of mean (SEM). Statistical analysis was carried out using PRISM 8 software-GraphPad.

## Data Availability

The authors declare that all data supporting the finding of this study are available within this article and its supplementary information files or from the lead corresponding author (G.P.L) upon request. The plasmids will be available from the lead corresponding author (G.P.L) upon request.

## Disclosure

All the authors declared no competing interests.

## Acknowledgements

This work was supported by the National Institute of Health (NIH) grants, R21EY025034 and R01EY030889 to G.P.L.; 2R01DK087956-06A1, R56-DK116887-01A1, and 1R03TR003038-01 to D.N.; EY027013-02 to M.R.B.; and R01EY027355 to S.H. This project was also supported in part by a DCI research grant (019898-001) and by a SCTR-NIH/NCATS grant (5UL1TR001450) to G.P.L. The pCDNA3 Rod Opsin construct was a gift from Dr. Robert Lucas (Addgene plasmid # 109361; http://n2t.net/addgene:109361; RRID:Addgene_109361). The authors thank George Robertson (Keyence Microscopes) for the use of the Keyence BZ-X800 scope for semi-thin plastic sections, H&E sections, and immunofluorescence imaging. The authors thank Dr. Don Rockey (MUSC) and Dr. Seok-Hyung Kim (MUSC) for recommending suitable liver function tests and/or for providing the ALT liver function kit. We also thank Dr. Linda McLoon (University of Minnesota) for critical review of the manuscript.

## Author Contributions

G.P.L. and D.N. designed the research studies and wrote the manuscript. G.P.L., D.N., B. Rohrer, M.R.B., S.H., H-J. K., R.M., and J.S. edited the manuscript. G.P.L., A.K.S., M.R.B., R.D.M., E.O., D.N., E.A., B.R., S.W., S.H., and R.A.N. conducted experiments and acquired data. A.K.S., G.P.L., M.R.B., S.W., S.H., J.S., R.D.M., R.A.N. and D.N., analyzed and interpreted the data. M.R.B. and S.H. performed ERG and interpreted the data. R.D.M and R.A.N. performed ECHO and/or interpreted the data. R.M., H-J. K., R.A.N., R.D.M., M.R.B., J.S., S.H., B. Rohrer and J.H.L., supplied reagents, software, or provided equipment for data analysis. All authors have read and agreed to the published version of the manuscript.

## Supplementary Figures

**Fig. S1:**
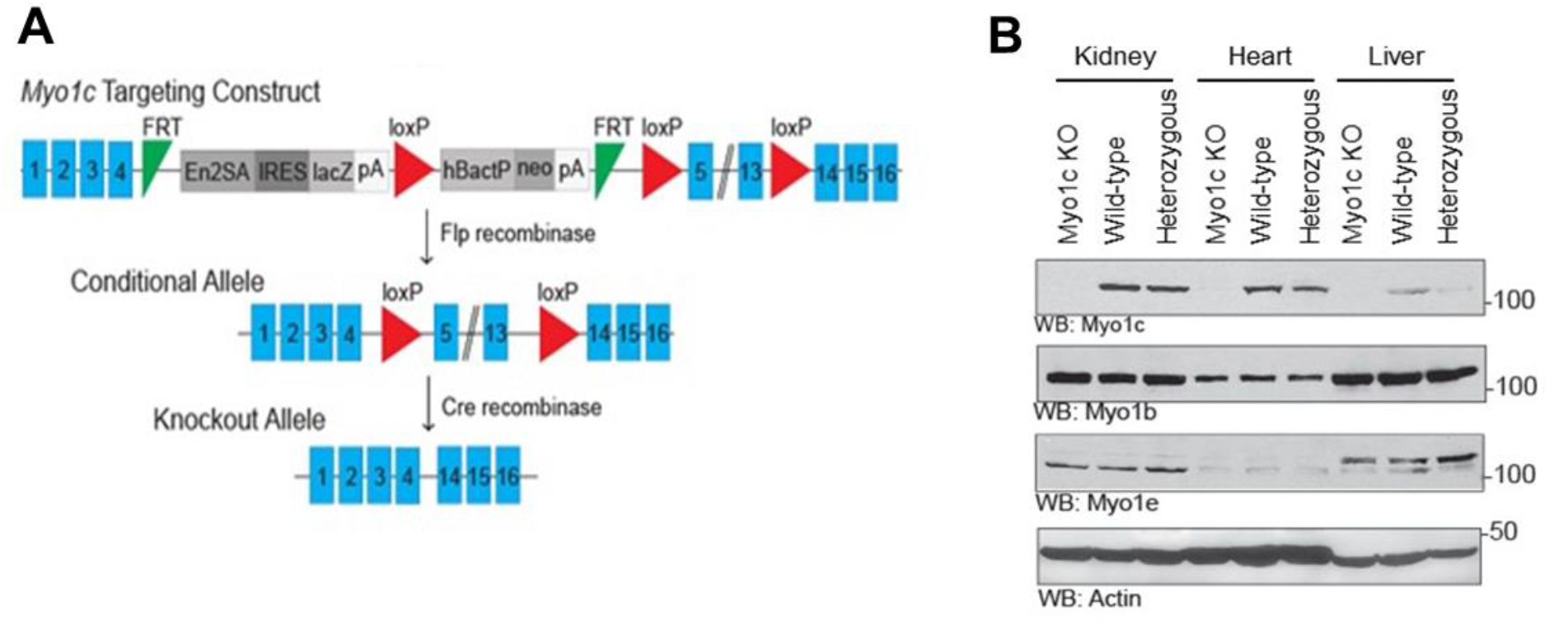
*Myo1c* targeting construct and Western blot analysis of MYO1C in systemic tissues: (**a**) Schematic representation of the Myo1c targeting construct for generation of the *Myo1c*-KO mouse line. (**b**) Western blot analysis confirmed MYO1C absence in various systemic tissues of *Myo1c-*KO mice. Absence of MYO1C did not affect MYO1B or MYO1E expression. Actin was used as the protein loading control. Representative images from multiple western blots (*n*=3) from *n*=3 animals per genotype.

**Fig. S2:**
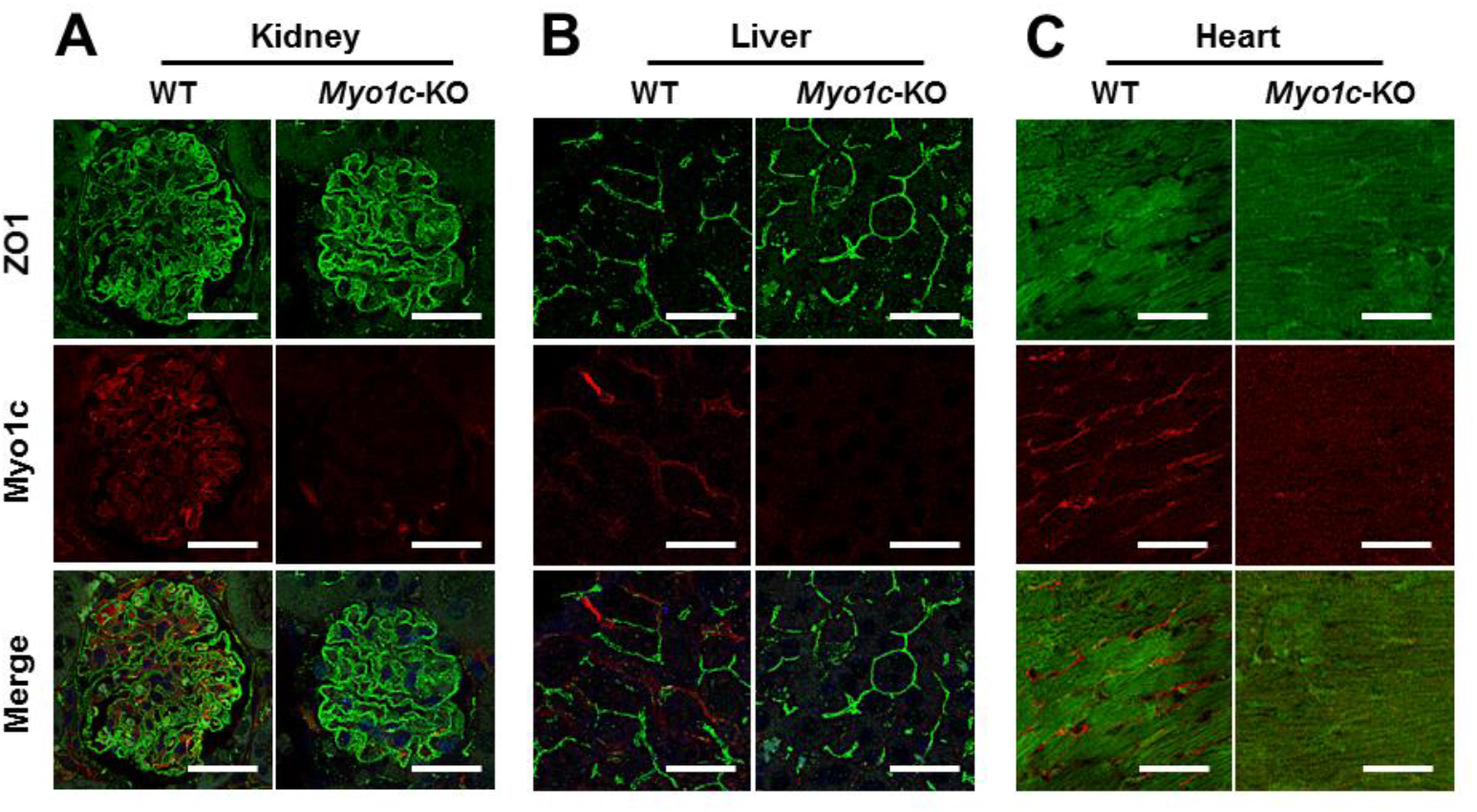
MYO1C expression in mouse tissue by immunofluorescence: Expression of MYO1C and ZO1 in systemic tissues, kidney (**a**), liver (**b**), and heart (**c**) of WT and *Myo1c*-KO animals (*n*=3 per genotype) by immunofluorescence. WT, wild type; KO, knockout. Representative images from *n*=3 animals. (**a, c**) Scale bar=50 μm; (**b**) Scale bar=75 μm

**Fig. S3:**
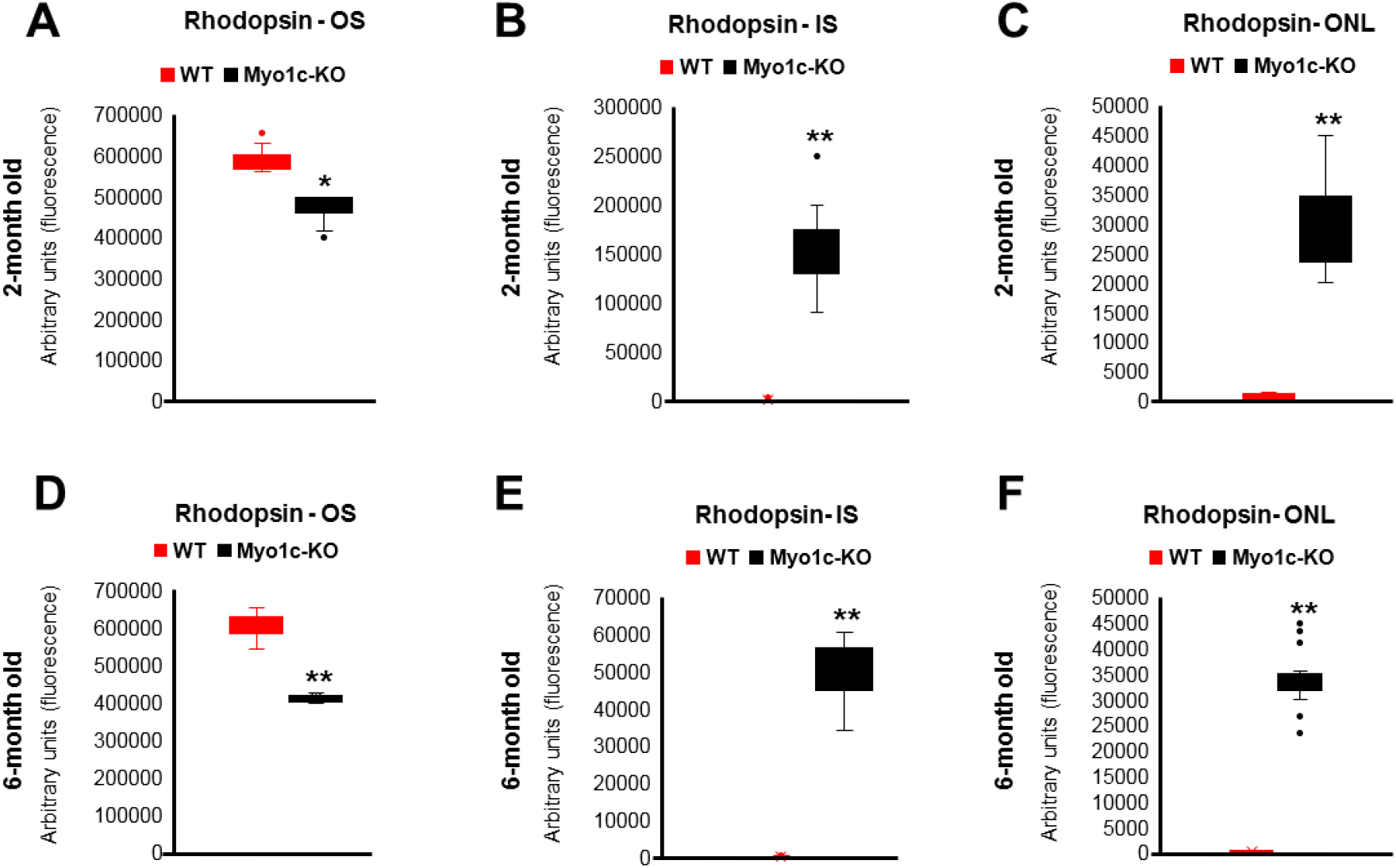
Quantification of OS lengths and distribution of Opsins in retinas of *Myo1c*-knockout mice from Figure 3: Rhodopsin distribution within rod OS, IS, and ONL were quantified in two month old animals (**a, b**, and **c**, respectively) and in six month old animals (**d, e**, and **f**, respectively). For quantification of rhodopsin distribution, 5-7 retinal sections from each eye (*n*=8 animals for each genotype and time-point) were analyzed using Image *J* or *FIJI* software. Mann-Whitney *U* test was used for statistical analysis and represented in Box-Whisker plots and considered significant ^*^*p*<0.05; ^**\*\***^*p*<0.005.

**Fig. S4:**
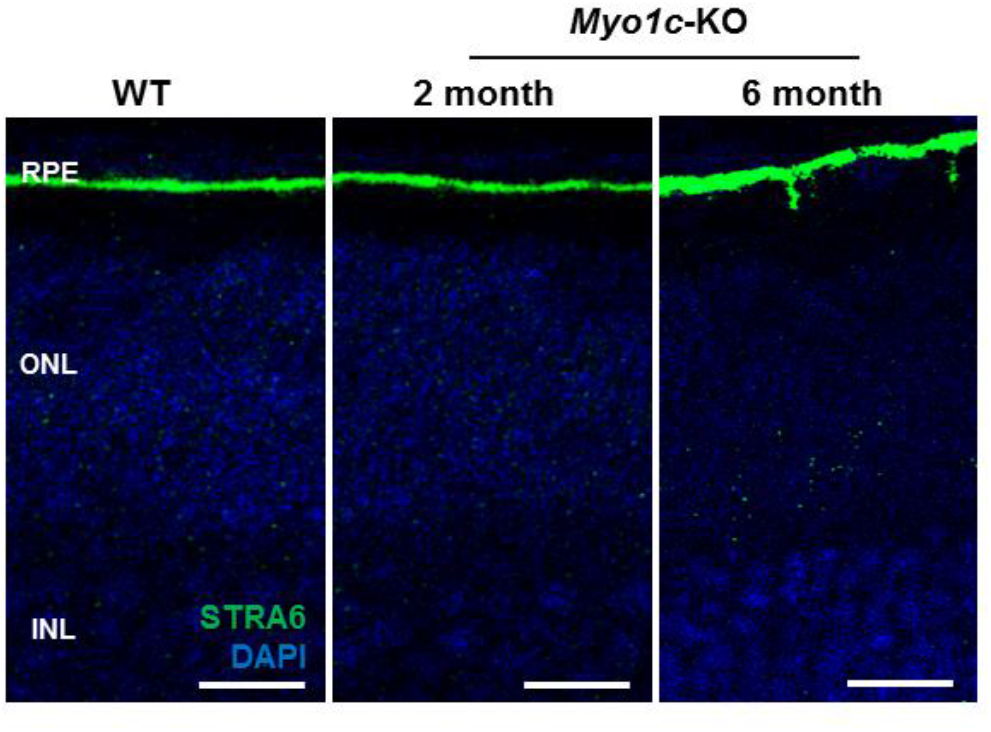
Immunohistochemical analysis of protein in RPE of wild-type/WT and *Myo1c*-knockout mice retinas: The RPE specific protein STRA6 was used to evaluate integrity of the RPE in both WT and *Myo1c*-KO mice. Representative images from *n*=8 animals (5-7 sections per eye) per genotype and age. Scale bar=50 μm. RPE, retinal pigmented epithelium; ONL, outer nuclear layer; INL, inner nuclear layer.

**Fig. S5:**
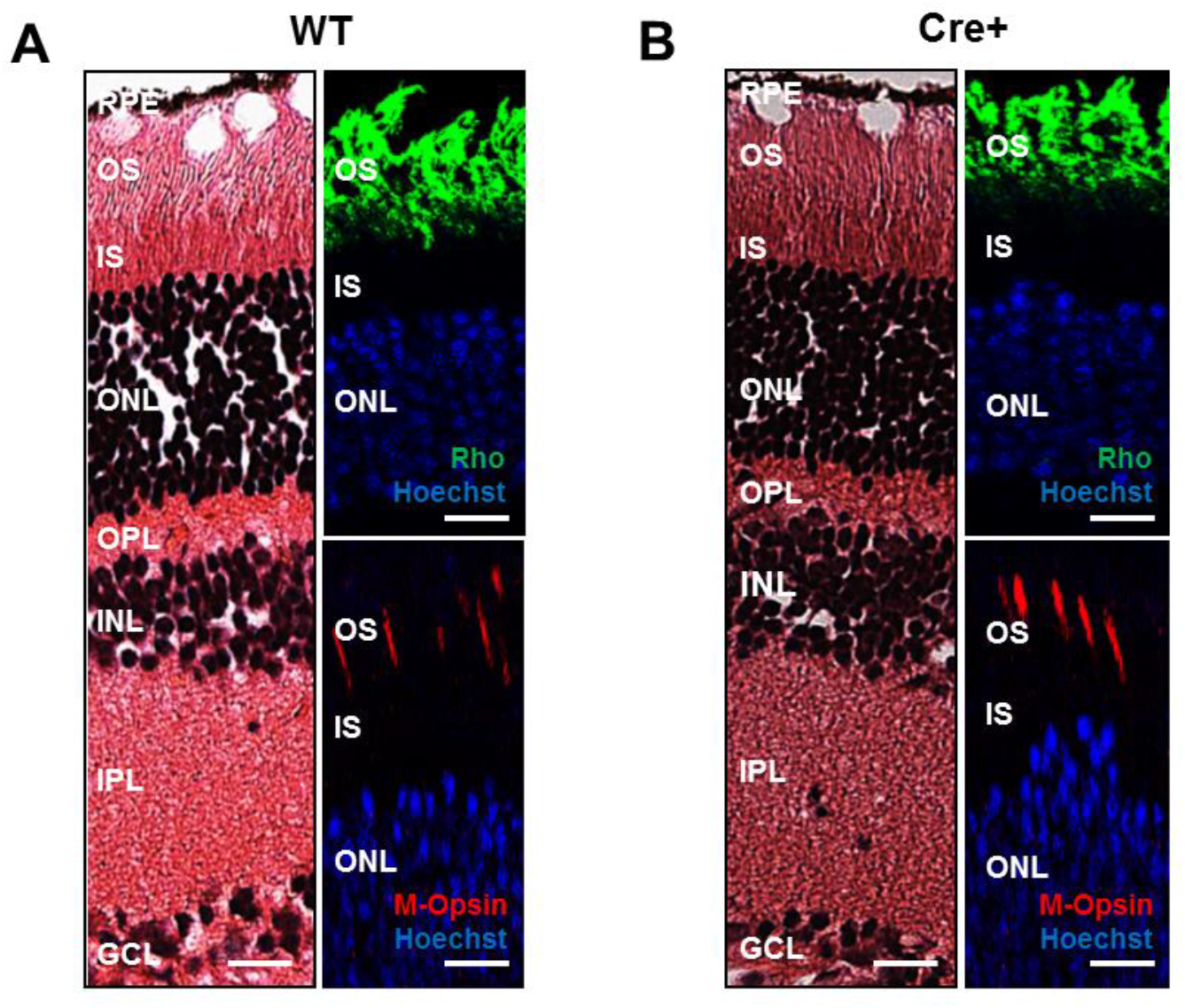
Immunohistochemical and histological analysis of retinas from Cre+ mice: Retinal histology, H&E staining, and localization of rhodopsin (Rho), red/green cone opsin (M-opsin) in WT (**a**) and Cre+ (**b**) mice. (**a, b**) Immunofluorescence Scale bar=50 µm. (**a, b**) Histology (H&E staining) Scale bar=100µm. Representative images from *n*=3 animals each genotype at 2-3 months of age (5-7 retinal sections per eye). RPE, retinal pigmented epithelium; OS, outer segments; IS, inner segments; ONL, outer nuclear layer; OPL, outer plexiform layer; INL, inner nuclear layer; IPL, inner plexiform layer; GCL, ganglion cell layer.

**Fig. S6:**
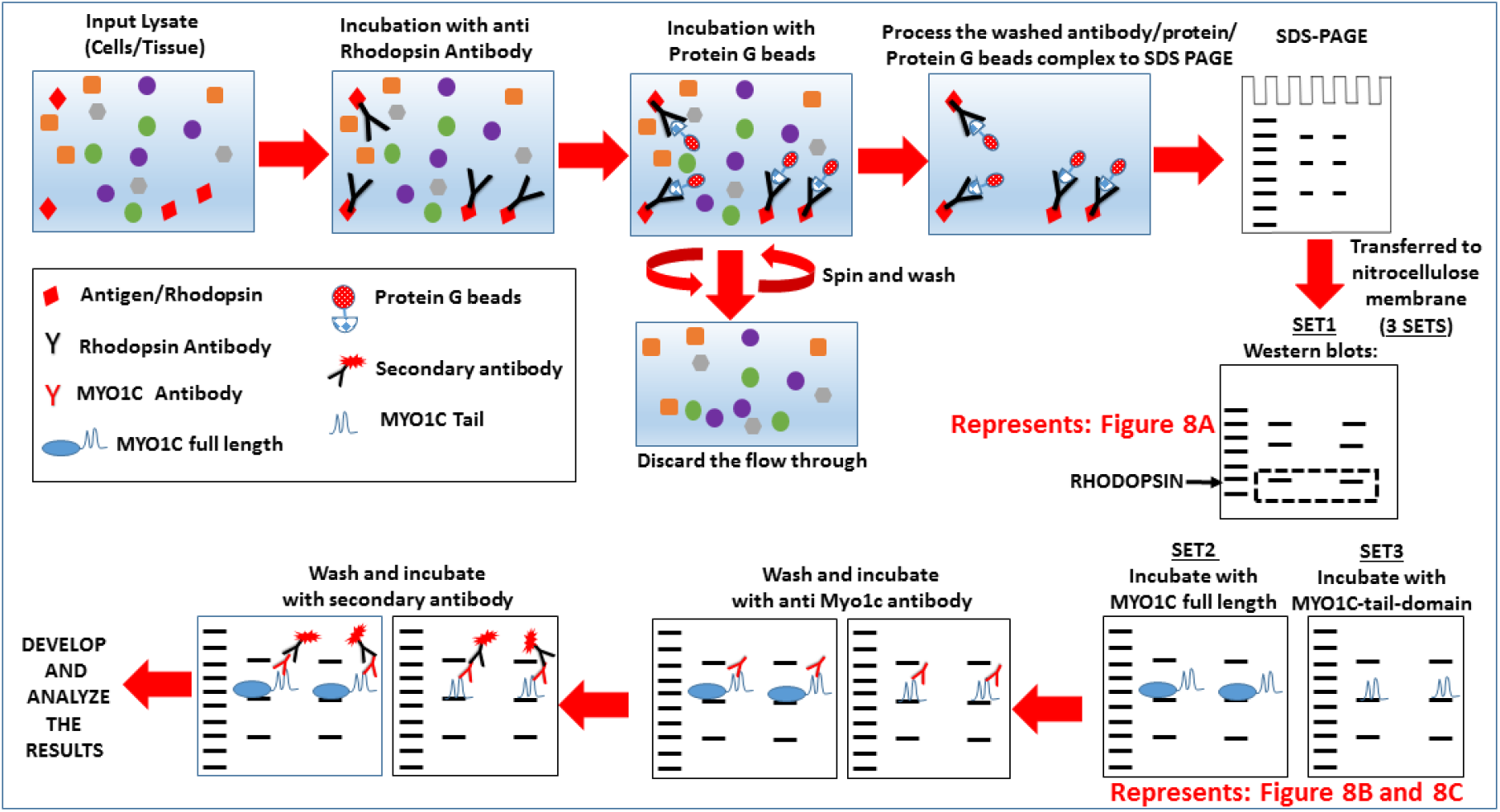
Schematic representation of MYO1C-Rhodopsin Co-IP binding experiments from figure 8: Detailed flow chart of Co-IP binding experiments representing figure 8 are shown.

**Fig. S7:**
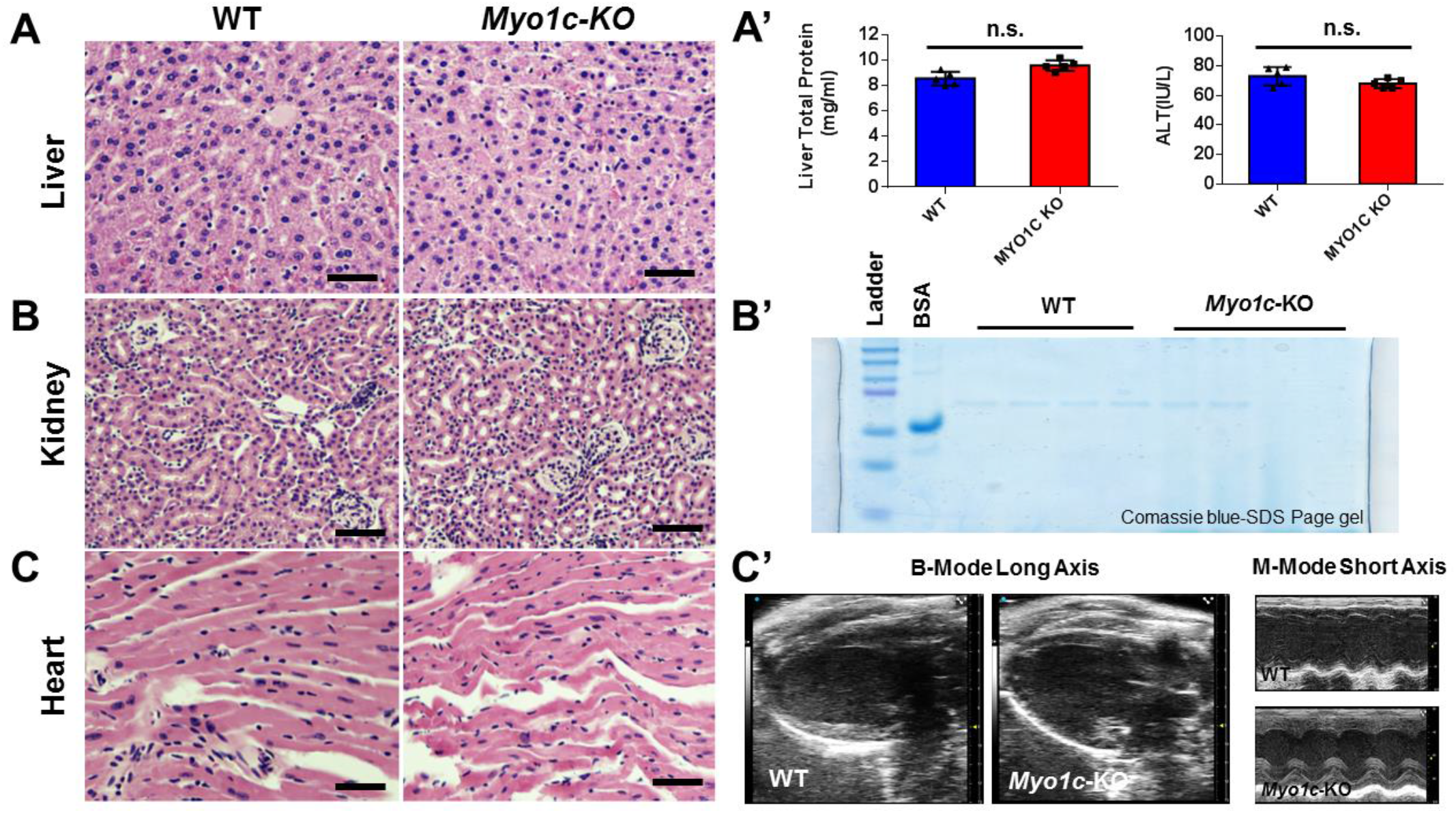
Representative tissue histology and functional analysis of systemic organs from WT and *Myo1c*-KO animals: Systemic organs, liver (**a**), kidney (**b**), and heart (**c**), of *Myo1c* and WT animals (*n*=4 each genotype at 3-4 months of age) were sectioned and stained with haematoxylin & eosin (H&E) to evaluate for systemic pathology of whole-body MYO1C loss. Functional analysis of systemic organs using ALT liver function tests (**a’**), urine analysis for kidney proteinuria/albuminuria (**b’**), and heart function using Echocardiography (**c’**) was performed in *Myo1c*-KO animals and compared to WT controls. **a’**, Liver function tests by Alanine Aminotransferase/ALT assay. No significant change (n.s.) was found in total protein concentration or ALT activity of liver (*p*>0.1) in *Myo1c*-KO compared to WT mice. Five biological replicates were used for each assay and statistical test used was one-tailed students *t*-test. **b’**, approximately 20µl of urine from *Myo1c*-KO and WT animals were electrophoresed on SDS-PAGE gels and stained with Comassie blue. 5ug/ml BSA was used as positive control. BSA, bovine serum albumin. **c**’, representative images from WT and *Myo1c*-KO heart showing B-Mode long axis and M-Mode short axis. (**a, b, c**) Scale bar=50µm.

**Fig. S8:**
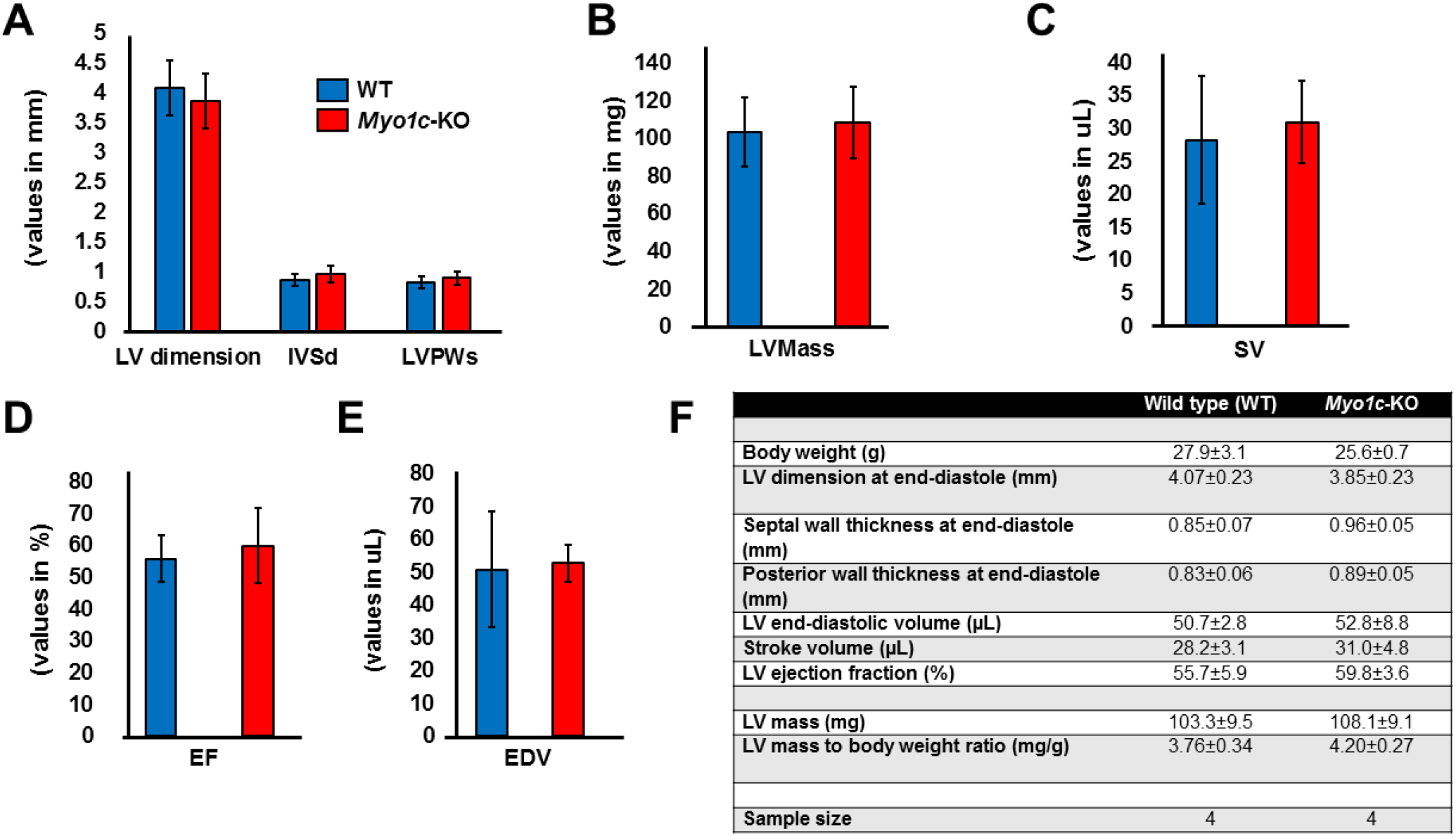
Detailed Echocardiographic (ECHO) parameters in WT and *Myo1c*-KO animals: Echocardiographic measurements was taken using the vevo 2100 ultrasound imaging system, to access cardiac function among genotypes. (**a**) Left-ventricle at end-diastole, Posterior wall thickness, and Septal wall thickness; (**b**) Left-ventricle (LV) Mass;Stroke volume (SV); (**d**) Left-ventricle ejection fraction (EF); (**e**) Left-ventricle end-diastolic volume (EDV); and (**f**) tabular summary of ECHO values from *n*=4 animals/genotype at 3-4 months of age. By *t*-test, there were no statistically significant differences between the two genotypes. Values presented as Mean±SEM.

**Fig. S9:**
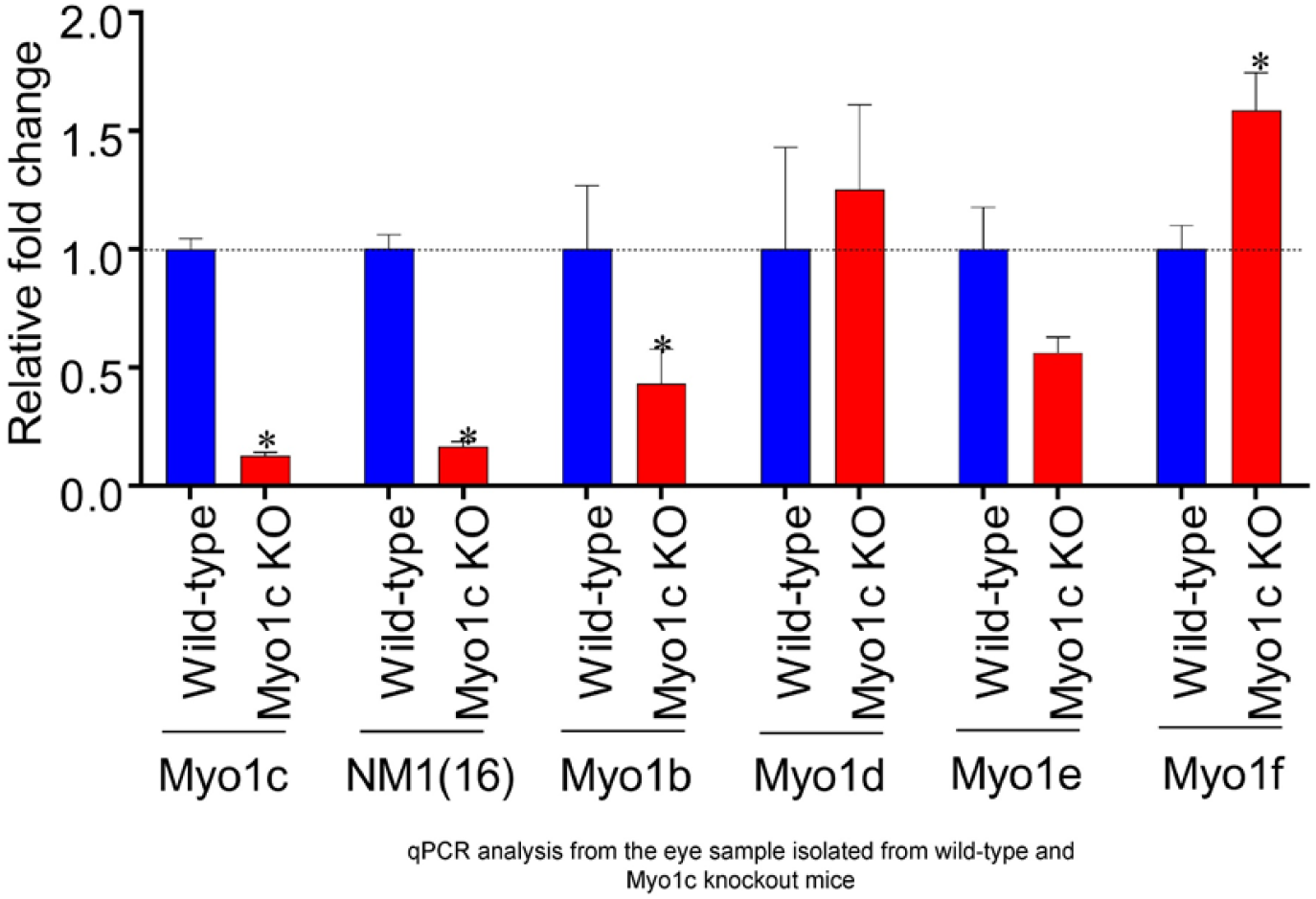
qPCR analysis of various MYO1C family members in mice retina: Retinas from WT and *Myo1c*-KO mice were isolated and processed for qPCR analysis using specific primers for various *Myo1c* family members including *Myo1b, d, e and f*. qPCR analysis was performed in triplicates for each sample and repeated thrice with freshly synthesized cDNA for each repeat experiment.

## Notes

### Competing Interest Statement

The authors have declared no competing interest.

### Summary of Updates

This version of the manuscript has been revised based on the reviewers comments. The revised version is improved to a significant extent for the better understanding of the readers.

